# Assessing Garmin’s Stress Level Score Against Heart Rate Variability Measurements

**DOI:** 10.1101/2025.01.06.630177

**Authors:** Hadar Rosenbach, Alon Itzkovith, Yori Gidron, Tom Schonberg

## Abstract

Daily stressors elicit physiological and mental responses impacting health, cognition, and behavior. Accurately assessing stress responses in natural settings remains challenging despite extensive research, though wrist-worn devices have the potential to address this gap through remote data collection. The Garmin fitness tracker provides a stress score largely based on HRV which must be validated prior to use in research. This study aimed to assess the stress score given by the Garmin Vivosmart 4 against HR and HRV from ECG recordings derived by the Polar H10 chest strap. A pilot study of 29 participants was conducted, followed by power calculations and preregistration of the main study which included 60 participants. Data were collected simultaneously from both devices during a laboratory session of restful and mental-stress-inducing tasks. Garmin’s stress score, mean HR, SD2/SD1, and HF power exhibited significant differences between stress and rest conditions. Moreover, Garmin’s stress score correlated significantly with HR, RMSSD, and SD2/SD1. Our findings suggest that physiological responses to mental stress were influenced by sex and tonic HRV. The study suggests that the GSS is indicative of mental stress, with its accessibility and noninvasive nature promising widespread utilization in various research domains.

## Introduction

In our daily lives, we encounter a multitude of triggers that elicit various forms of stress responses related to challenging or threatening situations [1]. These physiological and mental reactions play a pivotal role in shaping our health, cognition, and behavior [2], [3]. Mental stress has emerged as a pressing concern in contemporary society, exerting a profound impact on both physical and psychological well-being. The term ‘stress’ encompasses the body’s response to internal or external conditions (i.e., stressors) that disrupt homeostasis, triggering physiological and adverse psychological changes.

Stress involves the activation of two major pathways: the Sympathetic-Adrenal-Medullary (SAM) axis, which releases noradrenaline and instigates the “fight or flight” response, and the Hypothalamus–Pituitary–Adrenal (HPA) axis, responsible for the release of glucocorticoids, primarily cortisol in humans [4]. Physiological manifestations of stress include heightened sympathetic nervous system (SNS) activity and diminished parasympathetic nervous system (PNS) activity, resulting in increased heart rate, blood pressure, muscle tension, and cortisol levels, along with changes in blood flow and perspiration [5].

While the identification of stress biomarkers has been largely investigated in laboratory settings using physiological or psychosocial stressors [6]–[8], assessing stress in everyday life such as work-related pressures, social interactions, and environmental factors remains challenging [9]. This challenge is due to the complexity of real-world environments, individual responses, measurement inconsistency, and technological accuracy and reliability limitations [10].

Several physiological biomarkers have been used in the laboratory to identify stress, such as increased cortisol levels, which are detectable through saliva, blood, urine, and hair samples. This method typically requires multiple samples taken throughout the day along with a baseline measurement and necessitates laboratory equipment. Other methods including the analysis of salivary alpha-amylase (SAA), changes in heart rate, blood pressure, and the release of catecholamines that include epinephrine and norepinephrine, have been used in many studies as valid and reliable markers of the autonomic nervous system (ANS) activity and identification of stress [11].

Heart rate variability (HRV) reflects the fluctuation in time intervals between successive heartbeats, which is influenced by heart-brain interactions and ANS dynamics [12], [13]. It represents the ability of the heart to respond to physiological or environmental stimuli [14]. Moreover, HRV has emerged as a reliable biomarker for assessing autonomic nervous system (ANS) activity in a non-invasive and relatively simple manner [12]. Heart rate is regulated simultaneously by the two branches of the ANS; the parasympathetic nervous system (PNS) and the sympathetic nervous system (SNS) [13]. Changes in the activity levels of these branches are reflected through HRV and are associated with stress [4], [15]. Therefore, it is considered a reliable biomarker of ANS activity in response to stress [12].

The heart responds much faster to parasympathetic stimulation of the sinus node, which is the heart’s natural pacemaker responsible for generating and regulating its rhythmic contractions, compared to sympathetic stimulation. This dominance of the parasympathetic system causes higher frequency changes or higher variability in the heart rate signal. Thus, higher HRV is typically associated with parasympathetic activation, indicating a state of relaxation - “rest and digest” state, while lower HRV is linked to sympathetic dominance during “fight or flight” responses to stress or challenges and during physical exercise [14], [16], [17]. Conversely, In “fight or flight” responses, the sympathetic branch dominates, resulting in lower HRV [14], [16], [17].

The current evidence suggests that HRV is sensitive to induced mental stress in various methods and supports its use for an objective assessment of stress [12]. Neuroimaging studies suggest a link between HRV, and brain regions associated with reduced threat perception. HRV is seen as a mean to gauge the functional integration of brain regions involved in stress regulation and the flexible control over the ANS [12], [15]. This evidence further supports the utility of HRV as a reliable indicator of stress.

The term cardiac vagal tone reflects the PNS’s influence on heart regulation and is associated with various psychophysiological phenomena, including cognitive, emotional, social, and health-related self-regulation [18]. Consequently, psychophysiological research has focused specifically on HRV variables that assess vagal tone [18]. In addition, HRV also serves as a valuable predictor of risk for several chronic non-communicable diseases (NCDs) such as cardiovascular diseases, stroke, and cancer [19]. Research has shown that HRV, which reflects vagal nerve activity, provides insights into the role of autonomic nervous system in disease development [20]. Low HRV (or reduced vagal tone) has been linked to higher mortality and morbidity from NCDs, as it reflects dysregulated physiological processes such as excessive SNS activity, inflammation, and oxidative stress—key contributors to the pathophysiology of these diseases [21], [22].

HRV is represented by a set of statistical metrics derived from an RR signal (the time interval between consecutive R-wave peaks on an electrocardiogram) or inter-beat interval (IBI). HRV can be analyzed in the time-domain, in the frequency-domain and through non-linear indices [13].

Temporally, the main variable of interest is the standard deviation of the inter-beat-interval (IBI) of normal sinus beats (SDNN, i.e. the square root of variance, measured in ms)[23]. “Normal” means that abnormal beats, like ectopic beats (heartbeats originating outside the right atrium’s sinoatrial node), have been removed [13]. A higher SDNN typically signals healthy autonomic balance with strong vagal tone (PNS dominance), whereas a lower SDNN might suggest stress, reduced vagal tone, or an overactive SNS. SDNN is more accurate when calculated over 24 hours and is an accepted measurement of cardiac health [13]. The root mean square of successive differences between normal heartbeats (RMSSD) reflects vagal tone [24], [25] and is more influenced by the PNS than the SDNN [18]. The percentage of adjacent NN (normal-to-normal) intervals that differ from each other by more than 50 ms (pNN50) is also an accepted correlation with PNS activity [13], [17], [18].

In terms of frequency, signals are typically filtered into four bands: ultra-low frequencies (ULF, below 0.0033 Hz), very-low frequencies (VLF, between 0.0033 and 0.04 Hz), low frequencies (LF, between 0.04 and 0.15 Hz), and high frequencies (HF, between 0.15 and 0.40 Hz). While the High-frequency band reflects vagal tone [18], the low-frequency band reflects a mix of both the sympathetic and parasympathetic branches [4], [23], [26]. Frequency domain measures can be normalized to represent the relative value of each power component proportional to the total power minus the VLF component (LF [nu] = LF/(total power-VLF)×100, HF [nu] = HF/(total power-VLF)×100). High-frequency power is highly correlated with the pNN50 and RMSSD temporal measures [24]. Lower HF power is correlated with stress, panic, anxiety, or worry [24]. The ratio of LF [ms2] to HF [ms2] power (LF/HF ratio) was long considered to represent the sympatho-vagal balance [18]. However, this view has been criticized, today’s consensus is that the precise physiologic utility of the ratio is unclear [27], [28]. A loose relationship between LF power and sympathetic nerve activation is one of the main causes of ambiguity in the metric [27]. Furthermore, the SNS contribution to LF power varies profoundly with testing conditions such as anatomical position [13], [17]. Thus, the LF/HF ratio is now considered to have a low predictive value [18], [27].

In addition, non-linear indices could be obtained from the IBI signal. One of those non-linear indices is the Poincaré plot - a scatter plot in which every R–R interval is plotted against the prior interval. It illustrates patterns of HRV in the shape of an ellipse. Analysis of the Poincaré plot yields two indices resulting from the orthogonal distances between the scatter and the elliptical diameters. Crosswise, the standard deviation of the distance of each point from the y = x-axis (SD1), specifies the ellipse’s width and measures short-term HRV in milliseconds. Lengthwise, the standard deviation of each point from the y = x + average R–R interval (SD2) specifies the ellipse’s length. SD2 is affected by the overall variability of the signal and hence measures short- and long-term HRV in milliseconds [13], [18]. The ratio SD2/SD1 provides insights into the balance between short-term and long-term HRV. A higher SD2/SD1 ratio suggests more pronounced variability in longer-term heart rate patterns relative to short-term fluctuations, often associated with healthier autonomic nervous system function. In a previous study conducted by Melillo et al., nonlinear HRV metrics, including the SD2/SD1 ratio, were found to be effective in detecting real-life stress conditions, such as during a university examination [29].

Studies of HRV reactivity to psychological stressors in healthy human participants show that stress-induced changes in HRV metrics correlated with a reduction in vagal tone that was quantified by decreased levels of RMSSD, pNN50, HF, increased levels of SD2/SD1 and LF/HF, and an increase in HR [12], [18], [24], [30].

The recording period for HRV metrics has been standardized in 1996 by the Task Force of the European Society of Cardiology (ESC) and the North American Society of Pacing and Electrophysiology (NASPE) to ensure consistency across studies for the measurement, interpretation, and clinical use of HRV [23]. To standardize different studies investigating short-term HRV, a 5-minute recording was recommended unless the specific nature of the study required a different approach. A shorter recording could be performed; while the recording should last for at least 10 times the wavelength of the lower frequency bound of the investigated component with a minimum of approximately 1 minute to assess the HF components of HRV, approximately 2 minutes are needed to address the LF component [23].

Variables such as age, sex, and health status are crucial for interpreting HRV measurements, particularly for ultra-short-term (UST) recordings of less than 5 minutes [13]. Multiple studies have shown that temporal HRV measurements tend to decrease with age, with the most significant decrease between ages 20 and 30 [31]. Additionally, one study found [32] that in 24-hour recordings, a decline in SDNN between the ages of 40 and 100 was present and a U-shaped pattern of RMSSD and pNN50, declining from ages 40 to 60 and then increasing after age 70. A large meta-analysis [33] suggested that women have higher average heart rates and lower values for certain HRV measures, such as SDNN and SDNN index, particularly evident in studies spanning 24 hours. They also exhibited lower total, VLF, and LF power, but higher HF power compared to men. Despite their higher average heart rate, women tend to show relative dominance of the PNS, while men show relative dominance of the SNS. Furthermore, a study examining the correlation between HRV and habitual aerobic exercise [34], has shown that higher fitness levels were associated with significantly higher levels of HRV.

To measure HRV, a heart rate signal with high time resolution must be recorded. This can be achieved through several methods, primarily electrocardiogram (ECG) recording or photoplethysmography (PPG). The gold standard and most accurate technique is ECG recording [18]. ECG equipment records the heart’s electrical activity, allowing direct detection of the QRS complex. The QRS complex graphically represents the depolarization of the ventricles, indicating the moment when the ventricles contract to pump blood out of the heart [35]. The R wave, the largest deflection on an ECG signal, represents the main muscle contraction. By identifying the R wave on each heartbeat, the time interval between two consecutive heartbeats, known as the RR interval, can be measured.

In research settings, ECG is often performed using non-ambulatory recordings using multiple electrodes placed on the chest and limbs connected to a stationary ECG machine. This method provides highly accurate and detailed recordings but is less convenient for long-term monitoring. For ambulatory monitoring, ECG can be recorded using a chest strap sensor connected via Bluetooth to another device, such as a phone or computer. Devices like the Polar H10 heart monitor are well-validated against the gold standard and shown to be relatively accurate [36].

Recent technological advances in heart rate monitors have enabled the remote collection of physiological data in natural environments beyond traditional laboratory settings [37]. Sensors using Photoplethysmography (PPG) technology can record a relatively accurate heart rate signal and can be employed in wrist-worn monitors, such as smartwatches or fitness trackers, as well as on the earlobe or finger. A PPG device contains a light source and a photodetector. It sends light toward the capillaries, and the photodetector measures the reflected light from the tissue. The reflected light indicates blood volume in the vessel, forming the basis of the heartbeat signal. This results in an IBI signal, which provides the time in milliseconds between heartbeats [18], [38].

While the PPG sensor is considered to represent an accurate approximation of the IBI [39], it has some limitations. PPG measures a mixture of the IBI and pulse transit time, and therefore, it cannot detect changes in arterial elasticity that occur under stress, which can lead to changes in pulse transit time. Consequently, its accuracy under stress might be compromised [17], [40]. Additionally, the R spike of the QRS complex in ECG is easier to detect clearly, allowing for manual artifact correction on the reading [18]. Overall, ECG offers the highest accuracy, while PPG provides convenient, non-invasive monitoring suitable for various settings.

The introduction of PPG sensors facilitated the exploration of various behavioral, cognitive, and health phenomena. Previous studies showed that ultra-short-term analysis of heart rate and RR intervals could reliably monitor mental stress in field settings [41]. The use of wearable heart rate monitors, specifically wrist-worn devices, has increased in the last decade. Many wrist-worn devices offer individual health and wellness monitoring mainly based on heart rate measurements using PPG technology [42].

A considerable number of studies have been conducted in the last few years to validate heart rate measurements of commercially available wrist-worn devices from manufacturers such as Apple, Fitbit, Garmin, Polar, and Samsung [43]–[45] and presented a wide range of accuracies. Furthermore, recent research evaluated the validity of HRV measurements provided by wrist-worn PPG technology compared to measurements extracted from ECG signals [46]–[48]. Regardless, a considerable portion of commercially available devices to monitor and improve personal health, have not been formally validated through independent research [49].

The Garmin Vivosmart 4 fitness tracker (GV4) (Garmin Ltd., Olathe, Kansas) is a wrist-worn device equipped with a PPG sensor designed for monitoring physiological indicators such as heart rate, VO2 max, as well as monitoring sleep, stress levels, and activity tracking. It generates a stress level score ranging from 0 to 100 every three minutes, utilizing a non-invasive method largely based on HR and HRV indices [50]. Despite these features, the device does not provide access to raw IBI data or detailed HRV metrics.

Furthermore, while the stress level score calculated by Garmin’s device relies on HRV, its validity has yet to undergo independent validation. The challenge is compounded by the restricted access to raw data, including IBI signals, which are necessary for thoroughly assessing the accuracy of these scores. In fact, certain studies have indicated that while HRV measurements under rest conditions demonstrate satisfactory accuracy, their reliability diminishes in dynamic scenarios, such as during cognitive and emotional stress [51]. This highlights the pressing need for robust validation studies, especially if PPG-based stress detection is to be effectively applied in future field studies.

Therefore, the objective of the current study was to assess the stress level scores given by the wrist-worn fitness tracker GV4 (Garmin Ltd., Olathe, Kansas) in identifying a physiological response to mental stress. We aimed to compare and evaluate Garmin’s stress score data with HR and HRV metrics as references extracted from Electrocardiogram (ECG) recording using a validated heart rate monitor - Polar H10 chest strap (Polar Electro Oy, Kempele, Finland). We chose the GV4 for its lightweight design and minimal interference with participants’ daily lives, including during sleep. Additionally, it maintains participant privacy by not independently measuring or storing location data, as it requires a smartphone GPS for location tracking. The study was pre-registered on the Open Science Framework platform [52] based on a pilot sample and a power analysis. In our experiment, participants wore a Garmin Vivosmart 4 fitness tracker and a Polar H10 chest strap heart rate monitor simultaneously to record heart rate signals during a laboratory session. During that time, participants underwent tasks that were designed to elicit varying levels of stress. HRV metrics were calculated from the heart rate signal recorded using the Polar H10 and stress level scores provided by the Garmin Vivosmart 4 fitness tracker. Changes in GSS across stress/rest conditions were compared with the changes in HR and HRV metrics to assess validity.

## Methods

Pre-registration and Pilot sample: a pilot experiment was conducted to assess the experimental protocol’s feasibility and estimate the main experiment’s sample size. Data from twenty-nine participants was collected and analyzed. A power analysis was conducted to estimate the sample size required to detect significant differences in GSS, HR, and RMSSD. The analysis yielded a sample size of sixty participants for the main experiment with a statistical power of 0.80 and a significance level of 0.0125 (applying Bonferroni correction to account for multiple comparisons of HRV metrics p < 0.05/4 = 0.0125).

Following the analysis of the pilot data, minor modifications were made to the experimental tasks and data collection to enhance variable control and improve data quality in the main experiment. Task durations were reduced to minimize participant fatigue and discomfort. Participants also provided self-reported stress assessments after each task, and additional data on sleep, fitness, and lifestyle were collected (see details in the main experiment procedure description). The main study was preregistered in OSF, outlining hypotheses, study design, and analysis plan prior to data collection, based on the pilot sample. An exploratory analysis that was not included in the pre-registration examined the effect of individual differences (e.g., sex, age, exercise habits, tonic HRV) on HR, HRV, and GSS. This deviation was implemented to enhance variable control and data interpretation. See more detail in the data analysis subsection.

Methods of the main study are provided below, and detailed methods of the pilot study are available in Section A of the supplementary material.

*Data sharing*: Experimental data, analysis codes, and task codes are available through the Open Science Framework (OSF): https://osf.io/5ugj3/?view_only=ac84346ee27c432b900b9273d375447f

Pre-registration is available at the OSF: https://doi.org/10.17605/OSF.IO/GDR2N.

### A. Participants

Participant recruitment and inclusion in the main experiment: Initially, a total of 90 individuals volunteered to participate in the study. During the procedural phase, seven volunteers were unable to undergo the second session due to personal reasons. Consequently, 83 participants completed the experiment. Following data collection and initial processing, 23 participants were excluded from the analysis due to missing data, yielding a final valid sample size of 60. Participant exclusion primarily stemmed from technical issues encountered during data collection, such as Bluetooth disconnections between the Polar chest strap and the smartphone application, resulting in an incomplete signal necessary for the calculation of HRV metrics. Additionally, a subset of participants exhibited too many missing data points in the GSS data. Participants were deemed ineligible for analysis if there were no recorded stress points during the entirety of one of the tasks.

The valid 60 participants (40 females) had a mean age of 27 years (SD = 5.6 years; range: 19-39). Participants were University student volunteers from social media advertisements. All participants were healthy, had normal or corrected-to-normal vision, normal hearing, reported no neurological conditions, did not consume any psychiatric medications or drugs, and were not diagnosed with cardiovascular irregularities. Female participants declared that they were not pregnant. Ethical approval was obtained from the ethics committee of Tel Aviv University, and all participants provided informed consent. Participants were compensated with a payment of 40 NIS per hour with a bonus payment ranging from 0-20 NIS was awarded according to individual success in the stress task.

### B. Procedure

To explore the relationship between HRV parameters and stress level scores obtained from the GV4 fitness tracker, data were collected concurrently using the GV4 and a Polar H10 chest-strap heart rate monitor (Polar Electro Oy) during the main laboratory session. The Polar H10 served as a validated device for recording RR signals extracted from the QRS complex using electrocardiography (ECG) technology. Stress level scores came from the GV4 which uses a PPG sensor.

The procedure included a preparatory meeting and an experimental task session. During the preparatory meeting, participants were provided with a GV4 fitness tracker synchronized to a designated personal account in the Garmin Connect mobile application on their phones. Demographic information including age, sex, height, and weight was collected. Additional behavioral information including smoking habits, oral contraceptive use, habitual exercise (frequency and intensity), and habitual sleep routines were collected. This information was used to explore how personal differences influence the physiological metrics assessed. Inclusion criteria were verified and informed consent was obtained after explaining the experimental procedures. To ensure accurate identification of stressful moments, participants were instructed to wear the fitness tracker for one full day and night prior to the main laboratory session, as per the recommendations outlined in the GV4 manual. Participants were instructed to abstain from food, caffeine, and smoking for at least two hours before the experimental task session, as well as intense physical activity, drugs, and alcohol in the 24 hours prior to the experimental task session while maintaining their regular sleep routine.

Participants came back to the laboratory for the experimental task session the following day or later in the week. At the beginning of the session, the experimenter ensured that stress and heart rate data had been collected using the GV4 during the preceding day and night. This laboratory-based session comprised three consecutive tasks each lasting 15 minutes (versus 30 minutes in the pilot experiment): a resting phase (baseline), a stressful phase, and a relaxation phase (recovery), for a total duration of 45 minutes.

Throughout the session, participants remained seated in front of a computer in a quiet laboratory testing room, wearing both the GV4 fitness tracker and the Polar H10 chest strap. They were instructed to minimize movements during the tasks, while data were simultaneously collected from both devices. During the resting phase, participants were instructed to sit quietly without any external stimuli. In the stressful phase, participants engaged in a computerized mental arithmetic task modeled after the “Montreal Imaging Stress Task” (MIST), a protocol used to induce psychosocial stress in participants [53]. Before they began, participants were informed of a monetary incentive contingent upon their performance. During the relaxation phase, participants viewed a video featuring natural landscapes and listened to calming music. Furthermore, participants were asked to provide self-reported stress assessments. Participants verbally reported their subjective assessment of mental stress levels on a scale from 1 to 10 after completing each task. This information provided additional insight into participants’ real-time stress experiences throughout the three mental states.

In addition, following the completion of the three phases, participants completed a Perceived Stress Scale questionnaire (PSS-14) [54]. The PSS-14 focuses on how uncontrollable or overwhelming respondents found their lives in the recent past, typically within a one-month period. It contains 14 items that ask about thoughts and feelings experienced recently, providing a broader view of overall stress. A visual representation of the experimental protocol is presented in Figure 1

**Figure 1.**
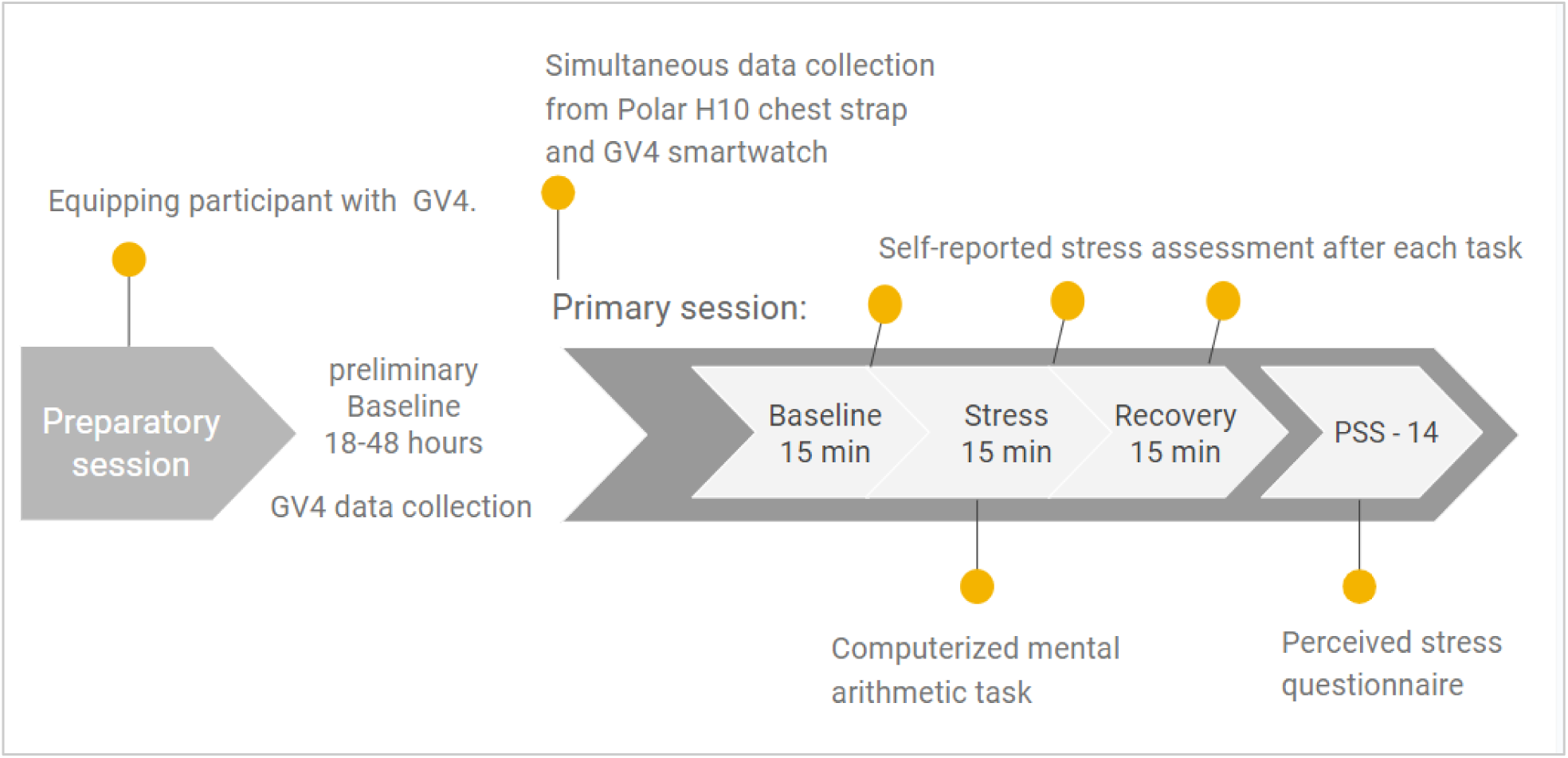
Protocol Flowchart of the Main Experiment.

**Figure 2.**
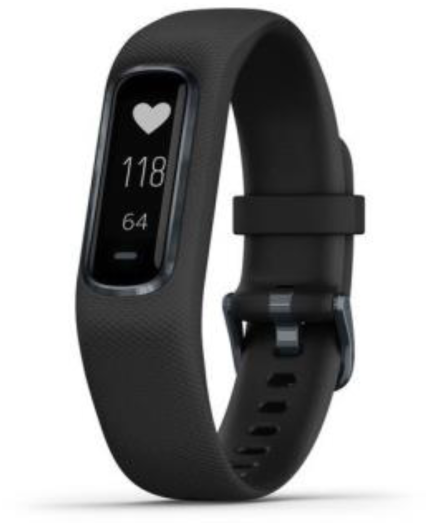
Garmin Vivosmart 4 fitness tracker. Source: Garmin’s official website

#### Data collection

the Garmin Connect mobile application (Garmin Ltd., Olathe, Kansas) was utilized to sync and store the stress data recorded by the GV4, using a designated account for each participant. Since direct access to the detailed data from the Garmin Connect app is restricted, we used Fitabase (Small Steps Labs, LLC), a specialized data management platform designed to integrate with the wearable device application. Fitabase provided an interface to securely access and download the stress data recorded by the Garmin smartwatches. All data retrieved through Fitabase was securely stored in a central database, ensuring participant confidentiality and data integrity. The RR signal was recorded using a Polar chest strap, with data captured and retrieved through Elite HRV installed on an iPhone13 Pro, a mobile application designed for HRV monitoring. The data was then downloaded from Elite HRV as a text file for further analysis, which involved calculating HRV metrics using Kubios HRV Premium software (version 3.5.0; Biosignal Analysis and Medical Imaging Group, Department of Physics, University of Kuopio, Kuopio, Finland) [55].

Reducing the duration of each task in the main experiment from 30 to 15 minutes relative to the task duration in the pilot procedure was implemented to minimize participant fatigue and discomfort, particularly during the stress-inducing task. Prolonged exposure to the stressor could lead to reduced engagement and potentially confound the physiological responses being measured. By shortening the task duration, we aimed to maintain participant compliance and data quality while still eliciting measurable stress responses. Additionally, shorter task durations allowed for increased experimental efficiency and participant engagement, facilitating data collection within a reasonable timeframe. However, the decision to limit task duration was constrained by the Garmin Vivosmart 4’s stress score readings. During the pilot study, a significant percentage of errors in the device’s readings led to missing stress score data. To ensure enough valid data points for analysis, we adjusted the recording length in the main experiment to allow between 1 to 5 stress score observations (each representing a 3-minute interval, totaling up to 15 minutes per experimental condition).

### C. Measures and Instruments

#### Wrist-Worn Fitness Tracker Garmin Vivosmart 4

The Garmin Vivosmart 4 fitness tracker (GV4) (Garmin Ltd., Olathe, Kansas) was utilized for monitoring physiological indicators including heart rate, VO2 max, alongside sleep patterns, stress levels, and activity levels. These measurements are collected using a wrist-worn PPG sensor. 2. depicts an image of the device. The GV4 provides a stress level score ranging from 0 to 100 at 3-minute intervals. This score is calculated by an algorithm developed by Firstbeat Technologies Ltd., a commercial company holding a U.S.-registered patent [51]. The algorithm detects stress through the segmentation and analysis of heartbeat signals. It utilizes HRV measurements derived from an IBI signal recorded by the PPG sensor. The stress-state detection procedure relies on two physiological assumptions [50]. First, it assesses sympathetic dominance relative to parasympathetic dominance, as inferred from heartbeat parameters. Second, it distinguishes the source of cardiac reactivity, excluding physical activity, movement, or posture. The required physiological information for state detection includes respiration rate, oxygen consumption (VO2), and excess post-exercise oxygen consumption (EPOC). These parameters are calculated within the Firstbeat algorithm based on HRV variables (time and frequency domain) and neural network modeling (Full product details appear [50]).

#### Heart Rate Monitor - Polar H10 Chest Strap

To obtain the RR signal for the calculation of HRV metrics we used the Polar H10 chest strap heart rate monitor (sampling rate: 1000 Hz; app software: Elite HRV App, Version 5.5.4) (Polar Electro Oy, Kempele, Finland) which has been validated against the ECG gold standard for heart rate and RR measurements [36], [56]. It has been previously used as a reference device for comparison in several studies [57]–[59].

Procedure: Participants wore the Polar H10 chest strap device, placed just below the chest muscles with wet electrodes for optimal conductivity. Recordings of RR intervals were sampled continuously during the experimental procedure, simultaneously with the collection of stress level score data with the GV4. The Elite HRV**©** mobile application was connected wirelessly through Bluetooth 4.0 signals coming from the chest strap to an iPhone13 Pro to record the RR signal. RR data was exported from the Elite HRV**©** mobile application to a designated email account.

#### Computation of Heart Rate Variability Metrics

Procedure: The raw RR data obtained by the Polar H10 strap were recorded using the Elite HRV app, exported as a text file, and analyzed with Kubios HRV Premium software (version 3.5.0; Biosignal Analysis and Medical Imaging Group, Department of Physics, University of Kuopio, Kuopio, Finland, [55], to obtain HRV parameters.

Prior to computing the HRV parameters, signal pre-processing was performed by the Kubios HRV Premium software, including noise detection, ectopic beats, and artifact correction using the built-in, automatic algorithm provided by the premium version of Kubios HRV. The correction was made by replacing identified artifacts with interpolated values using a cubic spline interpolation [55]. The automatic algorithm has been validated in previous work and found to be highly accurate [60].

To compare HRV values with corresponding GSS data given in 3-minute intervals, the IBI data were segmented into 3 minutes segments. HRV parameters were calculated within the time, frequency, and nonlinear domains from each 3-minute signal segment.

The spectral components were evaluated in fixed frequency bands, including very low frequency (VLF: 0–0.04 Hz), low frequency (LF: 0.04–0.15 Hz), high frequency (HF: 0.15–0.4 Hz), and low frequency to high-frequency ratio (LF/HF). A spectral analysis was performed using the autoregressive (AR) algorithm which has previously shown to generate a better representation of HF power by producing a spectrum with better resolution (for short excerpts) in comparison with the FFT method [18], [61].

Our investigation focused on HRV parameters reflecting vago-sympathetic balance and vagus nerve activity [18]. These key parameters include RMSSD (root mean square of successive differences between normal heartbeats) in the time domain, HF power (in normal units), and the LF/HF ratio (the ratio of low-frequency to high-frequency power) as an indicator of autonomic balance in the frequency domain. Additionally, the non-linear metric SD2/SD1 ratio (standard deviation of Poincaré plot lengthwise divided by the standard deviation of Poincaré plot crosswise) was included in our analysis [4], [18]. Based on existing literature, certain features were excluded from our analysis before pre-registration [18], [23]. Specifically, pNN50 and NN50 were removed due to their high correlation with RMSSD, with preference given to RMSSD. Additionally, TINN, HRV Tri Index, VLF, and log measurements were omitted as they were deemed more suitable for longer timeframes than those captured in our study. In summary, the pre-registered analysis included the HRV variables—RMSSD, HF power, LF/HF, and SD2/SD1—along with mean HR as variables of interest.

#### Mental Stress Task

A computerized task, based on the Montreal Imaging Stress Task (MIST), was designed to induce stress during the stressful segment of the three-task session. The MIST is a computerized protocol used to induce psychosocial stress in participants [53]. The protocol has two test conditions (control and experimental). In both control and experimental conditions, a series of simple arithmetic operations (sums and subtractions) were displayed on the computer screen which the participants should solve using the keyboard or the computer mouse. In the experimental condition, participants were under a time constraint that was meant to induce stress (see original paper for further details of this procedure [53]). Mental arithmetic tasks are a popular tool and are considered effective in mental stress induction for experimental purposes [62]. Specifically, the MIST protocol was used in several studies on stress responses and was found effective [63], [64].

In the present study, a similar protocol was used to induce stress among participants. A series of simple arithmetic operations (sums, subtractions, and multiplication) was presented on the screen and participants earned points for each correct answer they submitted depending on the response time. A faster response earned a higher score, and an incorrect answer reduced the score which discouraged guessing. A clock sound in the background was played to emulate the sense of time. Participant’s performance including their answers and response time were collected.

#### Perceived Stress Scale Questionnaire

The Perceived Stress Scale Questionnaire (PSS-14) [54] is a popular self-reported 14-item questionnaire for measuring psychological stress. It was designed to measure “the degree to which individuals evaluate situations in their lives as stressful” [54]. The PPS items are general and assess the feelings and thoughts of the individuals over the past month. Higher scores indicate higher self-reported perceived stress.

### D. Statistical Analyses

#### Main Experiment Pre-Registered Data Analysis

The first part of the statistical analysis of the main experiment was pre-registered prior to data collection, based on the analysis of the pilot study (details of the pilot study data analysis are available in section A in the supplementary material). It includes paired t-tests to compare HRV parameters, HR, and GSSs across baseline, stress, and recovery conditions to identify stress-related physiological responses to the stressful stimuli, as well an analysis to evaluate whether the stress score correlates with HR and HRV. The pre-registration did not include an exploratory analysis investigating the role of individual differences in stress modulation.

The dataset obtained in the main experiment consists of 60 participants, with each participant providing fifteen 3-minute interval observations. Each observation includes mean HR (bpm), HRV metrics (e.g., RMSSD [ms], HF power [nu]) derived from an IBI signal recorded by the Polar H10 chest strap, and stress scores from the Garmin fitness tracker. This resulted in a total of 45 minutes of data per participant, with each variable represented by a column in the dataset table.

The pre-registered analysis assessed the mean differences between stress conditions and both baseline and recovery phases, as well as computing the correlation between GSSs and selected HRV metrics. The selected metrics evaluated with GSS included mean HR (bpm), RMSSD (ms), SD2/SD1, HF power (nu), and LF/HF. Statistical significance was determined with p values < 0.05. To account for multiple comparisons (of HRV metrics), a Bonferroni correction was applied, setting the significance level to p < 0.05/4 = 0.0125.

We first identified stress-related physiological responses during the stress condition by comparing mean values of HRV parameters, HR, and GSSs across baseline, stress, and recovery conditions. Mean values were calculated for every metric in each condition, and paired t-tests were used to compare these values across the different conditions.

To examine the relationship between GSSs, HR, and HRV metrics, we employed two methods of correlational analysis. The first method involved a correlational analysis of condition means. For this analysis, we calculated the mean GSSs and HRV metrics within each experimental condition (baseline, stress, and recovery) for each participant. Using these means, we computed the Pearson correlation coefficient for each condition based on the data from all 60 participants. This method allowed us to assess the strength and direction of the relationship between HR, HRV parameters, and GSSs across different states. The second method was a within-subject analysis. For each participant, we calculated the Pearson correlation coefficient between GSSs and each HRV metric. We then averaged these individual correlation coefficients across all participants to obtain a mean correlation coefficient for each HRV metric. To determine the significance of these correlations, we conducted a one-sample t-test on the mean correlation coefficients. This process resulted in a table displaying the mean Pearson’s r and associated p-values for each HRV metric with GSSs, indicating whether there was a meaningful relationship between GSSs and HRV metrics across the participant sample. This two-pronged approach provided a comprehensive evaluation of the association between GSSs and HRV metrics, both at the group level and within individual participants.

#### Outlier Exclusion

Outlier handling was conducted post-registration to ensure the reliability and validity of the dataset. This procedure was performed in two stages. First, participants with HR exceeding 100 beats per minute (bpm) at rest during the baseline phase were excluded from the analysis, as such values are physiologically atypical. Second, for each metric within each experimental condition (baseline, stress, and recovery), the mean and standard deviation were computed. Observations exceeding ±3 standard deviations from the condition-specific mean were identified as outliers and excluded.

Following the removal of outliers, paired t-tests and correlation analyses were repeated using the cleaned dataset to assess the sensitivity of the findings to the presence of extreme values. Specifically, we re-examined whether the significance of variations in HRV metrics across experimental conditions changed and whether the strength of the relationships between the GSS and HR/HRV metrics was altered. This comparison allowed for an evaluation of the robustness of our results to outlier handling.

#### Exploratory Analysis: The Role of Individual Differences in Modulating HR, HRV, and GSS

Following the pre-registered analysis, we explored the effect of individual differences (e.g., sex, age, exercise habits) on HR, HRV, and GSS in stress reactivity. Linear mixed-effects models (LMM) were developed to assess the impact of physiological and demographic factors on GSS across different experimental conditions (baseline, stress, and recovery). The models incorporated independent variables such as sex, age, body mass index (BMI) in kilograms per square meter, regular exercise habits (at least 1-2 times a week or none), average nightly sleep duration in hours, oral contraceptive use, and baseline HRV category (low or high RMSSD). Two groups of HRV category were defined based on a median split on resting RMSSD (high versus low RMSSD taken during baseline condition). Additionally, interaction effects between sex and stress condition, and between HRV category and stress condition were investigated to test if the effect of the stress task differs based on sex or if individuals with low HRV versus high at baseline respond differently to the stress exposure. This type of model allowed us to capture both the fixed effects of the experimental conditions and other covariates, as well as the random effects due to individual variability. A model was built for each physiological metric of interest as the dependent variable: mean HR (bpm), RMSSD (ms), SD2/SD1, HF power (nu), and LF/HF. The analysis was conducted using the mixed linear model regression approach utilizing the ‘statsmodels’ package on Python. Significance was determined at p < 0.05, and coefficients with their respective p-values were examined to ascertain the strength and direction of the associations.

## Results

The following analyses were pre-registered prior to the commencement of data collection of the main study. Results of the pilot experiment are available in Section B of the supplementary material.

Data from 60 participants (40 females), with a mean age of 27.5 years (SD = 5.6), were valid for analysis. Participant’s demographic information and other lifestyle characteristics are provided in Table I.

**Table I:** Participant’s demographic information and lifestyle characteristics – Main Experiment.

| Numeric Variables |  |  |  |  |
| --- | --- | --- | --- | --- |
|  | Mean | Std | Min | Max |
| Age [years] | 27.5 | 5.6 | 19 | 39 |
| Height [cm] | 167.9 | 9.8 | 150 | 190 |
| Weight [kg] | 68.0 | 15.9 | 45 | 110 |
| Typical Sleep Duration [hours] | 7.1 | 1.3 | 5.0 | 9.0 |
| PSS-14 Score | 26.2 | 8.8 | 11 | 46 |
| Categorical Variables |  |  |  |  |

|  | Count | Percentage |
| --- | --- | --- |
| **Gender** |  |  |
| Female | 40 | 66.66% |
| Male | 20 | 33.34% |
| **Smoking** |  |  |
| Yes | 3 | 5.00% |
| No | 57 | 95.00% |
| **Oral Contraceptive Intake** |  |  |
| Yes | 33 | 55.00%<br>(82.50% of females) |
| No | 27 | 45.00%<br>(17.50% of females) |
| Exercise Regularly | 37 | 61.67% |
| **Exercise Frequency (times per week)** |  |  |
| Low (1-2) | 17 | 28.33% |
| Medium (3-4) | 15 | 25.00% |
| High (5 or more) | 5 | 8.33% |
| **Exercise Intensity** |  |  |
| Low | 6 | 10.00% |
| Medium | 21 | 35.00% |
| High | 10 | 16.67% |

### A. Variation of Measures Across Experimental Conditions

Stress Task Effects: Paired t-tests assessed significant differences in participants’ physiological measures across the three conditions (Baseline, Stress, and Recovery). A significance threshold was set at p < 0.0125 after applying the Bonferroni correction to account for multiple comparisons of HRV metrics.

The results of this analysis revealed significant differences in Mean HR between Stress vs. Baseline (p < 0.001) and Stress vs Recovery (p < 0.001), indicating elevated HR during stress-inducing tasks compared to rest periods. No significant differences were observed in the comparison of HR during Recovery vs Baseline. Mean RMSSD decreased during the stress condition relative to baseline and increased during recovery, however, this difference did not meet the significant threshold. The SD2/SD1 Ratio exhibits significant difference between Stress and Baseline (p = 0.0014). HF Power (nu) decreased significantly during the stress task compared to baseline (p = 0.003). LF/HF increased during the stress condition relative to baseline, but this difference did not meet the significant threshold. The GSS showed a very similar pattern to HR with a significant increase during the stress task (p < 0.001) and a significant decrease during recovery (p < 0.001). These findings underscore the dynamic physiological responses to stress-inducing tasks and subsequent recovery periods. See Figure 3 for a visual presentation of the comparison. For full results, please refer to section C of the supplementary material: Table S IV.

**Figure 3.**
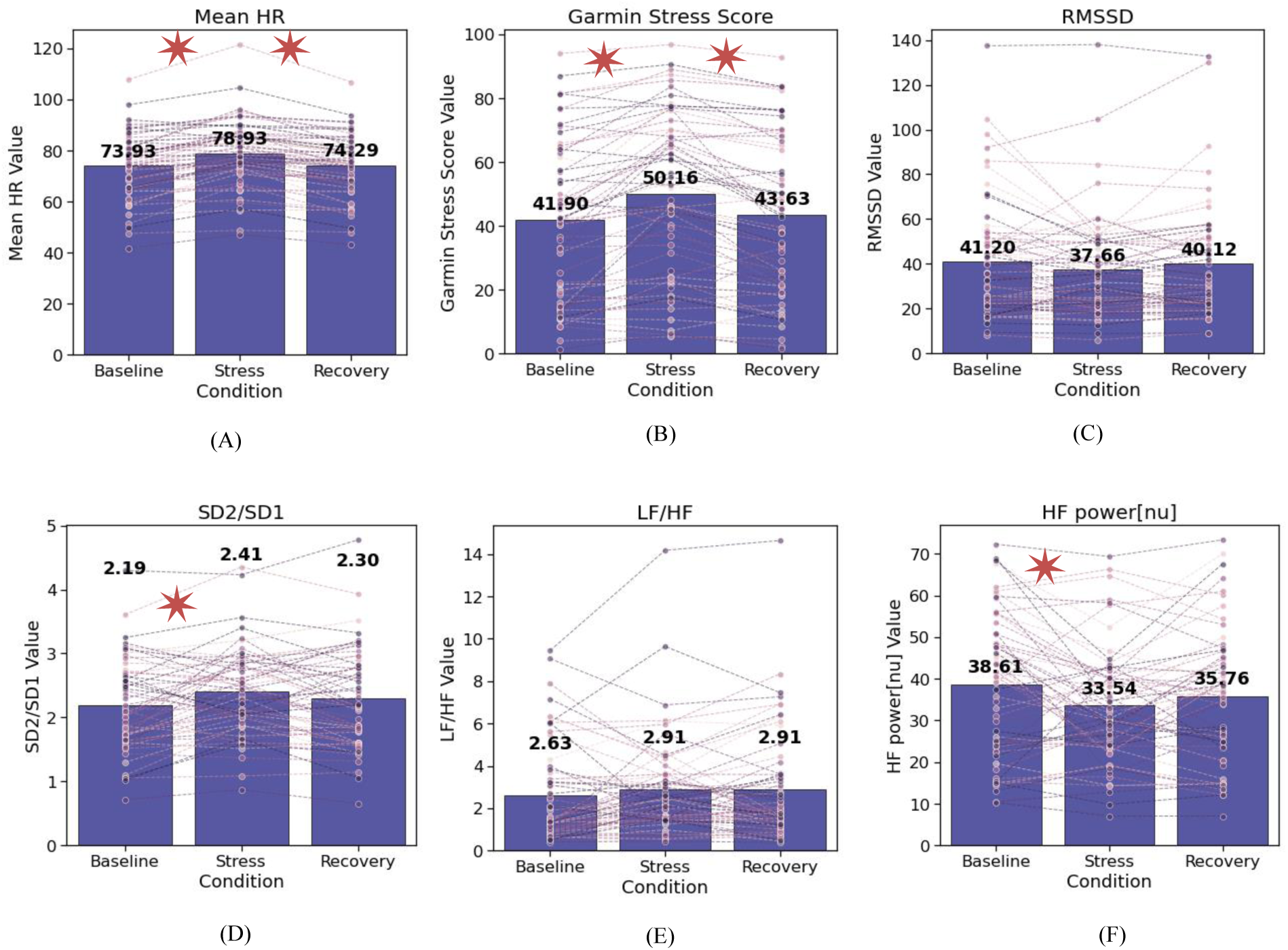
The distribution of participants’ HR, HRV, and GSSs across three conditions: baseline (rest), stress, and recovery. Notably, mean values in each experimental condition are indicated within the boxplot. Significant differences (P value < 0.0125) across conditions are marked by a star.

Alongside physiological measures, a paired t-test was conducted to examine differences in subjective self-reported stress levels (see Figure 4). On average, participants reported higher levels of subjective stress following the stress task compared to both baseline and recovery conditions (p< 0.001).

**Figure 4.**
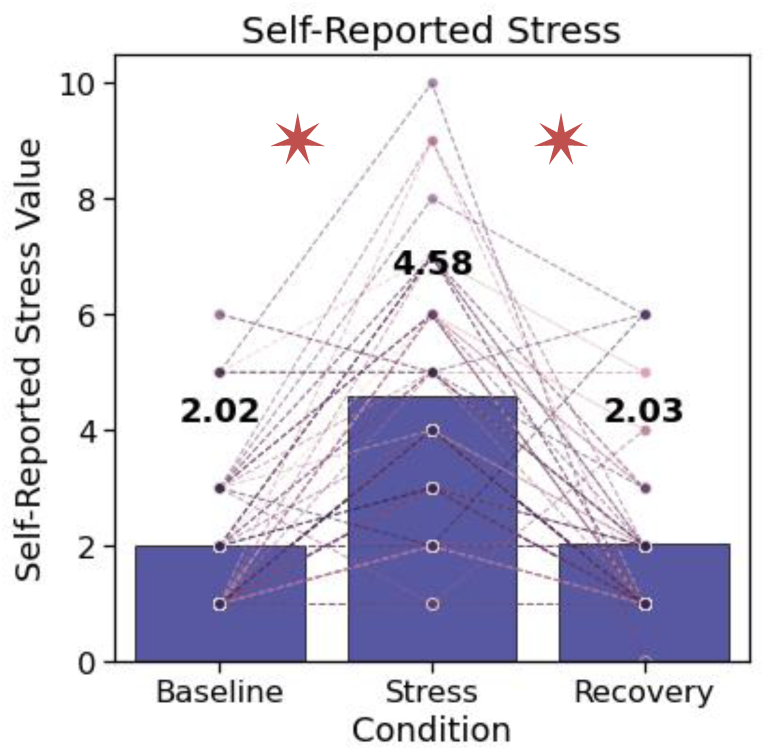
Subjective Self-Report of Stress Levels Across Conditions. Significant difference (P-value < 0.0125) across conditions is marked by a star.

### B. Correlation Analysis

Correlation analysis results revealed consistent associations between the GSSs and various physiological metrics, employing two different methods of computation.

#### Within-Condition Correlation

In Method 1, Pearson correlation coefficients were computed from mean scores across participants, within each condition (each participant had one observation in each condition, totaling 3 observations for participant). Robust positive correlations emerged between GSSs and Mean HR, as well as SD2/SD1, with coefficients ranging from 0.84 to 0.85 (p < 0.0001) and 0.61 to 0.64 (p < 0.0001), respectively. Conversely, a notable negative correlation was observed between RMSSD and Garmin’s metric, with coefficients ranging from −0.59 to −0.63 (p < 0.0001). Furthermore, lower positive correlations were found between GSSs and LF/HF (0.32 (p = 0.0126) to 0.38 (p = 0.0028)) and moderate negative correlations with HF (−0.4 (p = 0.0014) to −0.43 (p = 0.0006)). Comprehensive analysis data, including Pearson correlation coefficients and corresponding p-values for each metric across all conditions, can be found in section C of the supplementary material: Table S V.

#### Within-Subject Correlation

In Method 2, we investigated the relationships between GSSs and the physiological parameters at the individual level. Individual correlations per participant were computed, and the averages across participants were analyzed. This approach yielded a significant positive mean correlation between GSS and Mean HR (r = 0.74, SD = 0.35, p < 0.0001), indicating a consistent association between higher stress level scores provided by Garmin and elevated HRs. RMSSD displayed a lower negative correlation (r = −0.41, SD = 0.42, p < 0.0001), suggesting that increased GSSs were associated with reduced HRV. SD2/SD1 demonstrated a lower positive correlation (r = 0.32, SD = 0.45, p < 0.0001), indicating a tendency for higher GSSs to coincide with higher SD2/SD1 ratios. LF/HF and HF power exhibited low correlations with Pearson’s r of 0.18 (SD = 0.41, p = 0.001) and −0.20 (SD = 0.42, p = 0.001), suggesting weaker associations between Garmin’s metric and these metrics. High standard deviations were observed in the correlation statistics, indicating variability in these measures among participants. These findings highlight the complex interplay between physiological responses and stress levels, as captured by GSSs. A summary table of the results can be found in section C of the supplementary material: Table S VI.

#### PSS-14 Correlations

The analysis examined correlations between various physiological and self-reported measures and the PSS-14, a widely used questionnaire that assesses individuals’ perceived stress levels over the past month. Results revealed no significant relationships between the measures and PSS-14 scores. Pearson’s correlations were weak and non-significant: HR (r = −0.03, p = 0.797), RMSSD (r = −0.04, p = 0.769), SD2/SD1 (r = −0.06, p = 0.632), LF/HF (r = −0.07, p = 0.624), HF power[nu] (r = 0.01, p = 0.919), GSS (r = −0.17, p = 0.201), and self-reported stress (r = 0.25, p = 0.055).

Similarly, the stress to baseline differences showed weak and non-significant correlations: HR (r = −0.15, p = 0.266), RMSSD (r = 0.16, p = 0.229), SD2/SD1 (r = −0.11, p = 0.414), LF/HF (r = −0.13, p = 0.321), HF power[nu] (r = 0.04, p = 0.791), GSS (r = −0.19, p = 0.146), and self-reported stress (r = 0.11, p = 0.394). These findings suggest no meaningful relationship between the examined metrics and perceived stress levels as measured by the PSS-14 in this dataset.

#### Arithmetic Task Score Correlations

We further investigated the relationship between participants’ performance in the arithmetic task conducted during the stressful phase and their physiological metrics, as well as the disparity in physiological measures between conditions. The analysis revealed no significant relationships. Pearson’s correlations for baseline measures were weak: HR (r = −0.06, p = 0.670), RMSSD (r = 0.05, p = 0.720), SD2/SD1 (r = −0.00, p = 0.984), LF/HF (r = 0.09, p = 0.475), HF power[nu] (r = 0.04, p = 0.764), Garmin stress (r = −0.05, p = 0.684), and reported stress (r = −0.19, p = 0.140).

Similarly, the correlations for the stress to baseline differences were also non-significant: HR (r = −0.03, p = 0.829), RMSSD (r = 0.00, p = 0.979), SD2/SD1 (r = −0.06, p = 0.637), LF/HF (r = 0.06, p = 0.644), HF power[nu] (r = −0.04, p = 0.792), Garmin stress (r = 0.20, p = 0.131), and reported stress (r = −0.09, p = 0.486). These findings suggest no meaningful correlation between the performance in the arithmetic task and the physiological measures or the change in it across stress/rest state.

### C. Outlier Exclusion

One participant was excluded due to a resting heart rate exceeding 100 bpm, and 57 observations out of a total of 900 (15 per participant) were identified as outliers. These outliers, determined based on a statistical threshold of being 3 standard deviations or more from the mean, were identified separately for each experimental condition.

The removal of outliers led to marginal changes in the mean and standard deviation across conditions but did not alter the significance of the findings. Paired t-tests performed on the cleaned dataset continued to show significant variation in HR, SD2/SD1, HFnu, GSS, and self-reported stress across experimental conditions, consistent with the results of the original dataset. Similarly, correlation analyses revealed minor changes in Pearson r coefficients, differing by 0.01–0.04, while all correlations remained statistically significant. Across participants within each condition, the correlation between GSS and HR remained strong (Pearson r=0.84, p < 0.0001), while correlations with RMSSD and SD2/SD1 remained moderate (Pearson r=0.6±0.03, p < 0.0001). Correlations with frequency domain metrics (HFnu and LF/HF) remained weaker (with LF/HF Pearson r=0.3, p = 0.019, and with HF Pearson r=0.4, p = 0.014). Within-subject analyses yielded similarly minor differences in Pearson r coefficients (0.01–0.02). Full results of paired t-test and correlation analysis on the cleaned dataset are available in section C in the supplementary materials: Table S VII, Table S IIIII, and Table S IX.

### D. Exploratory Analysis: The Role of Individual Differences in Modulating HR, HRV, and GSS

We followed up the pre-registered analysis with an exploratory analysis to examine whether certain individual differences modulate the physiological measures we assessed. We tested the effect of sex, age, BMI, habitual exercise, average nightly sleep duration, oral contraceptive use, and baseline HRV (low vs. high based on a median split of resting RMSSD) on GSS, HR, HRV, and self-reported stress across the three conditions (baseline, stress, and recovery) through linear mixed-effects models (LMM).

The model included two main factors: sex (male vs. female) and baseline HRV category (low vs. high RMSSD). Both factors were allowed to interact with the experimental conditions to explore their combined influence on the dependent variable. To adjust for potential confounders, age, BMI, oral contraceptive use, and sleep duration were included as covariates. Additionally, a random intercept was incorporated for each participant to account for within-subject variability across repeated measures. Separate linear mixed-effects models were developed for each dependent variable: mean HR, HRV metrics, and self-reported stress. For example, formula (1) represents the model built to predict GSS:

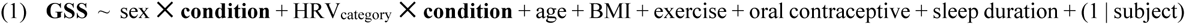

The split into two groups, low versus high baseline HRV, was performed by applying a median split of resting RMSSD values (median = 35.75 ms, during baseline condition). These groups, representing 30 participants each, differed significantly in RMSSD (Low HRV group: 22.2 ± 6.6 ms; High HRV group: 60.2 ± 23.6 ms, t[58] = 8.4, p <0.001).

#### Stress Task Effects

The LMM model revealed significant differences in the physiological metrics between conditions. Specifically, higher HR and SD2/SD1 values were observed during the stress task compared to baseline (β = 6.305, p < 0.001 and β = 0.370, p < 0.001, respectively), while RMSSD was lower (β = −7.943, p = 0.001). In the frequency domain, HF levels decreased during the stress task compared to baseline (β = −8.505, p < 0.001) and, although they increased during recovery, they remained lower than baseline (β = −5.028, p = 0.023). The LF/HF ratio increase during the stress task compared to baseline was statistically insignificant.

##### GSS

A significant increase in GSS was observed during the stress task (β = 10.384, p < 0.001), with a marginally higher GSS also noted during recovery (β = 3.908, p = 0.048), both compared to baseline.

#### Self-reported stress levels

participants reported higher stress levels at the end of the stress condition compared to baseline (β = 2.764, p < 0.001), emphasizing the stress-induced task efficacy.

#### Individual Characteristics Modulating GSS

The LMM results revealed several significant predictors of GSS, including HRV category, exercise, BMI, and sleep duration. Participants with low baseline HRV exhibited significantly higher GSS compared to those with high HRV (β = 21.679, p < 0.001). Regular exercise and longer sleep durations were associated with lower stress scores captured by the GSS (β = −17.948, p = 0.002 and β = −4.304, p = 0.027 respectively). Higher BMI was associated with higher stress scores (β = 1.365, p = 0.018). No significant effects related to sex, oral contraceptive use, or age were found in Garmin stress scores. Interactions between sex and experimental conditions, as well as HRV category and experimental conditions, are not significant. Figure 5 demonstrates GSS across experimental conditions, categorized by individual differences such as Sex, Habitual Exercise, and Baseline HRV Category.

**Figure 5.**
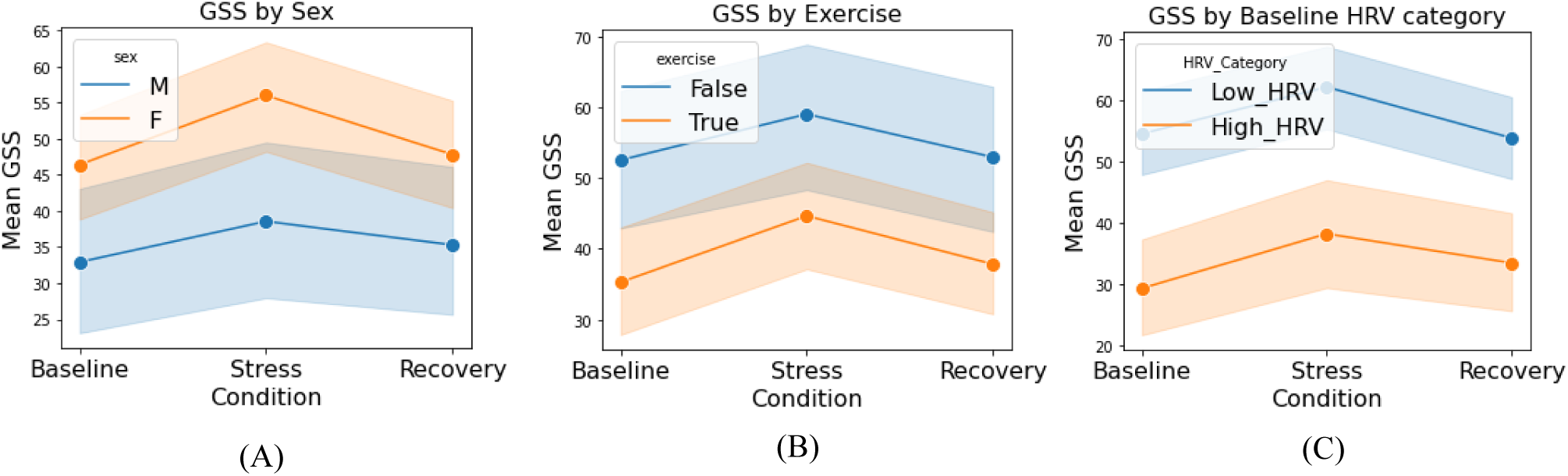
GSS Across Experimental Conditions, Divided by Individual Differences including sex (Female vs. Male), exercise regularly (‘True’ for at least 1-2 times a week vs. ‘False’ for none), and HRV category (low vs. high baseline RMSSD).

#### Individual Characteristics Modulating HR

The factors identified to influence HR significantly include baseline HRV category and habitual exercise. Specifically, participants categorized with low HRV exhibited higher HR compared to those with high HRV (β = 12.057, p < 0.001). Additionally, individuals who exercise regularly (n=37) have, on average, a lower HR (β = −9.830, p = 0.001) compared to those who do not exercise (n=23), see Figure 6. The within-subject random effect was significant (β = 0.158, p = 0.004), indicating variability in HR across participants.

**Figure 6.**
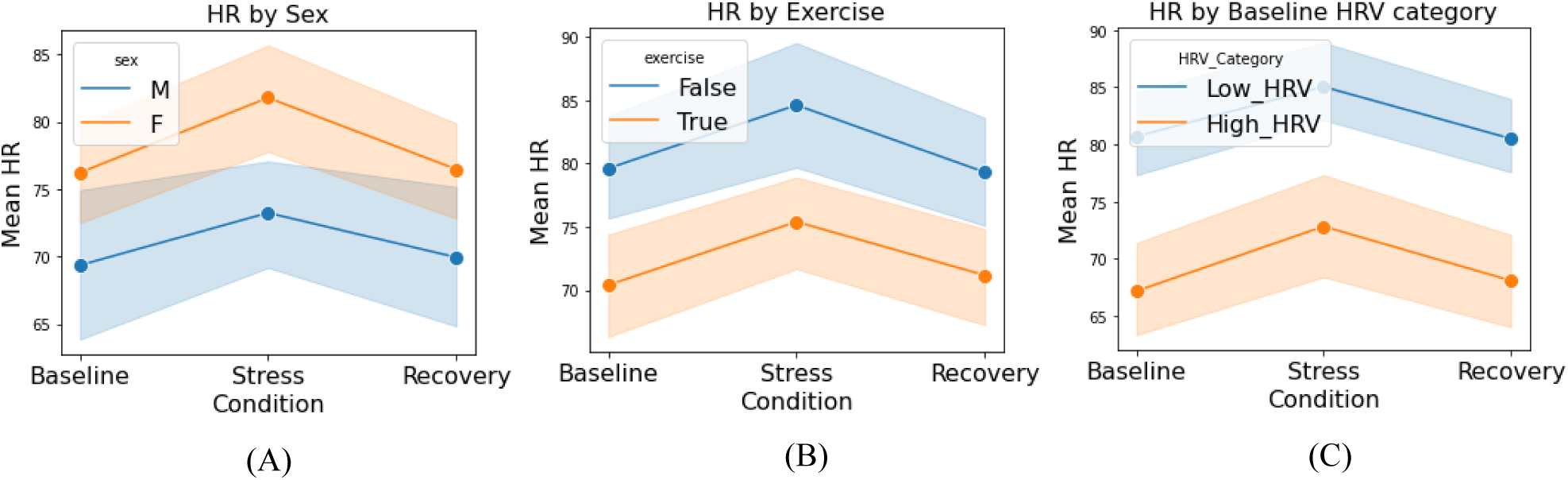
Mean HR Across Experimental Conditions, Divided by Individual Differences including sex (Female vs. Male), exercise regularly (‘True’ for at least 1-2 times a week vs. ‘False’ for none regularly), and HRV category (low vs. high baseline RMSSD).

#### Individual Characteristics Modulating HRV metrics

The mixed linear model analysis revealed that baseline HRV category influenced all the HRV variables. Sex influenced the frequency domain metrics. The interaction effect of sex with experimental condition modulates SD2/SD1 ratio and HFnu.

##### HRV category

As expected, the analysis indicates that participants categorized with low baseline HRV demonstrated a significantly lower RMSSD (β = −36.442, p < 0.001) and lower levels of HF (nu) (β = −17.621, p < 0.001). while low baseline HRV was significantly correlated with higher levels of SD2/SD1 ratio (β = 0.843, p < 0.001) and increased LF/HF ratio (β = 2.076, p < 0.001). See Figure 7 for a visual presentation of the differences.

**Figure 7.**
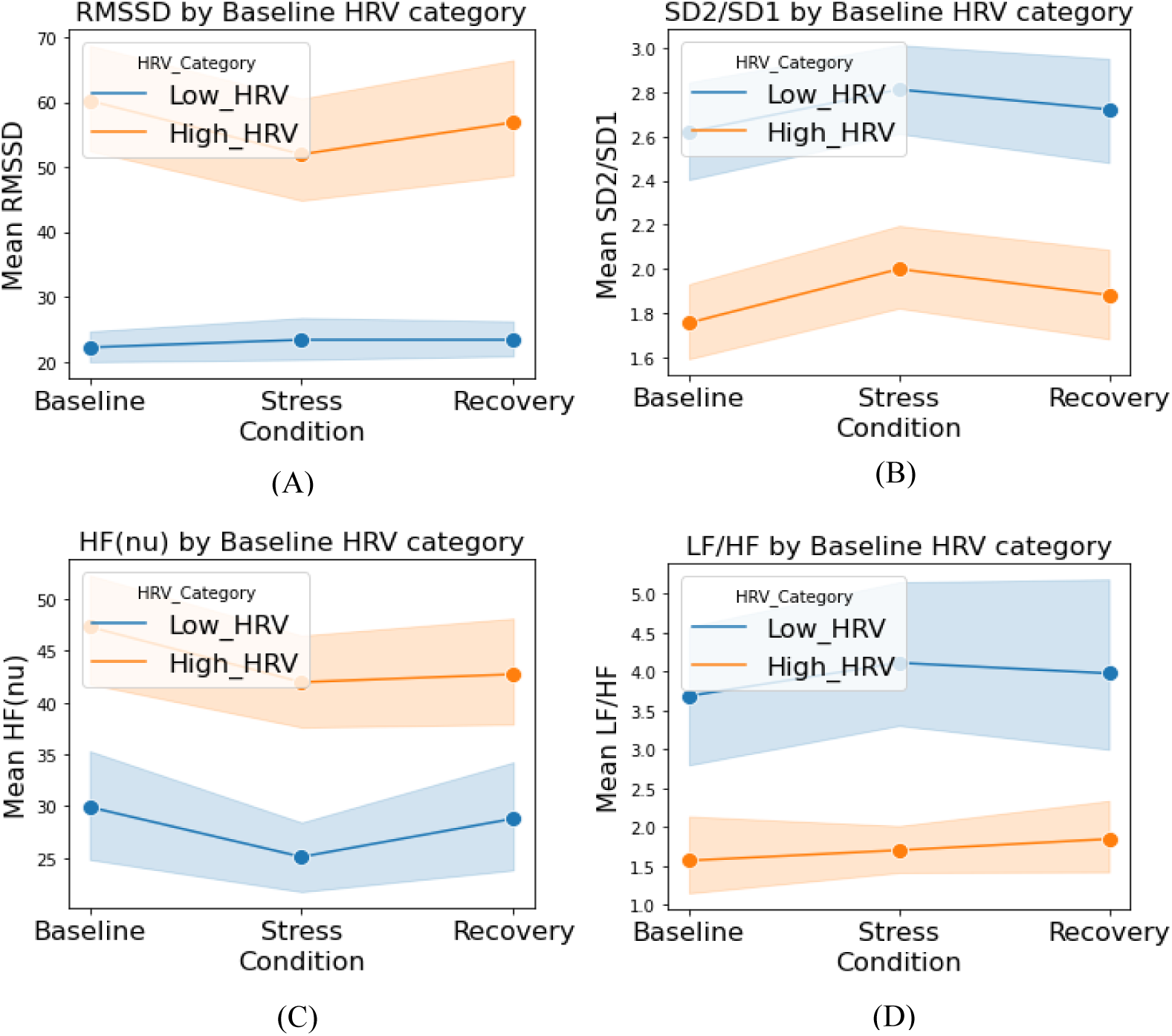
HRV metrics Across Experimental Conditions, divided by HRV category (low vs. high baseline RMSSD)

A significant interaction effect of the HRV category with the stress condition in modulating RMSSD was identified. The two groups - low vs. high baseline HRV display an opposite trend of RMSSD values across conditions (as shown in part A of Figure 7). Participants with high baseline HRV (n = 30) displayed a decrease in RMSSD (approximately 8.2 points) during the stress task, followed by a slight increase during recovery. While participants with low baseline HRV (n = 30) exhibited a marginal increase during the stress task (approximately 1.1 points). Yet, the within-group differences did not reach statistical significance.

##### Sex

An interaction effect of sex and the stress condition on SD2/SD1 ratio was significant with males showing a reduction in the SD2/SD1 ratio under stress compared to females who on average show increased values during the stress task (β = −0.344, p = 0.003). Sex had a significant effect on the frequency domain metrics: LF/HF ratio and HFnu. Males had higher levels of LF/HF (β = 1.772, p = 0.008), and lower levels of HFnu (β = −14.586, p = 0.001). A significant interaction between sex and condition suggests that the relative change in HFnu from rest to stress task is influenced by sex, with males experiencing an increase in HFnu during stress compared to females who experiencing a decrease (β = 8.661, p = 0.003). See Figure 8 for a visual representation.

**Figure 8.**
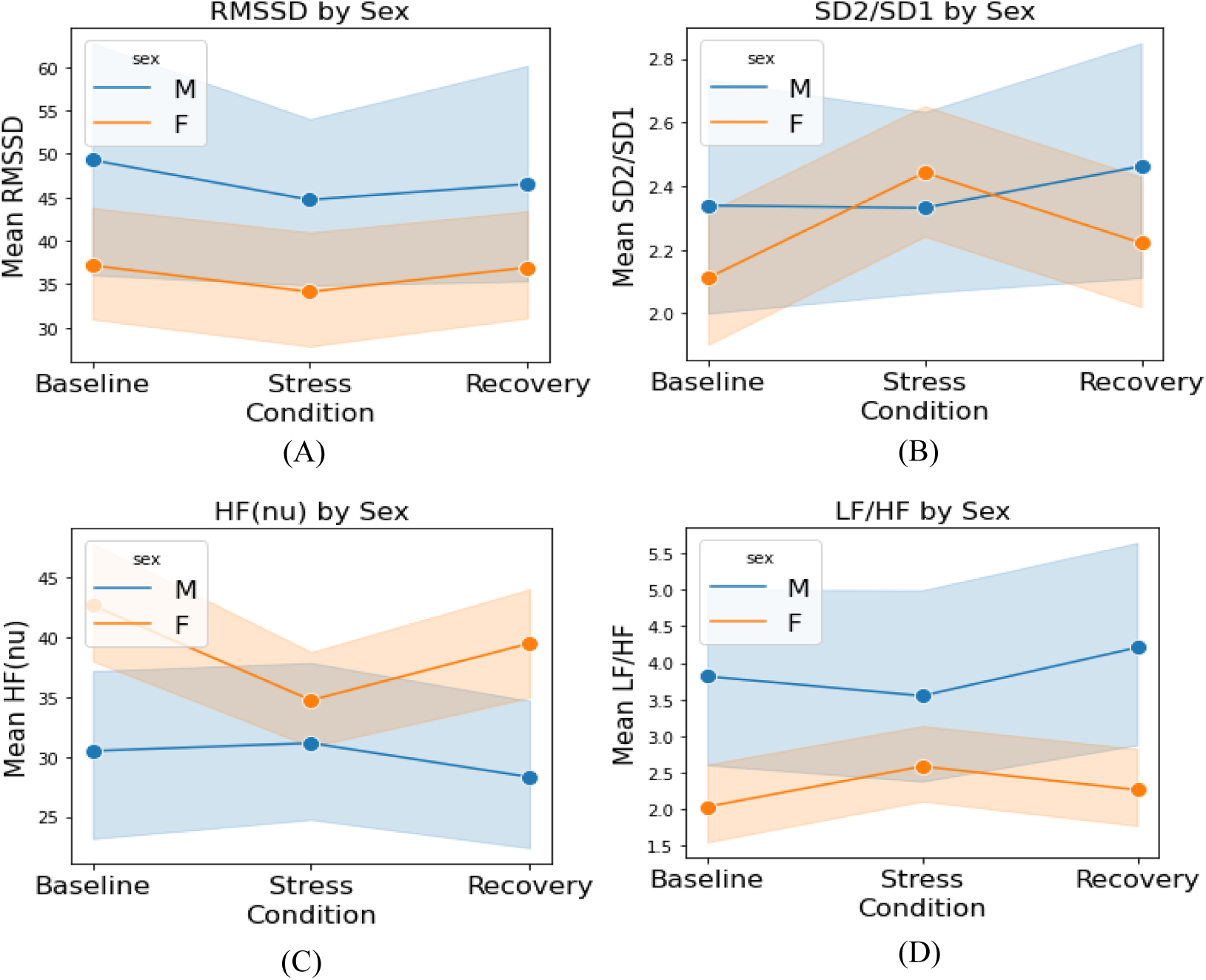
HRV metrics Across Experimental Conditions, divided by Sex (Female vs. Male).

Furthermore, we found a marginal positive effect of exercising regularly on RMSSD, however, it did not reach statistical significance (β = 9.771, p = 0.069). See Figure 9.

**Figure 9.**
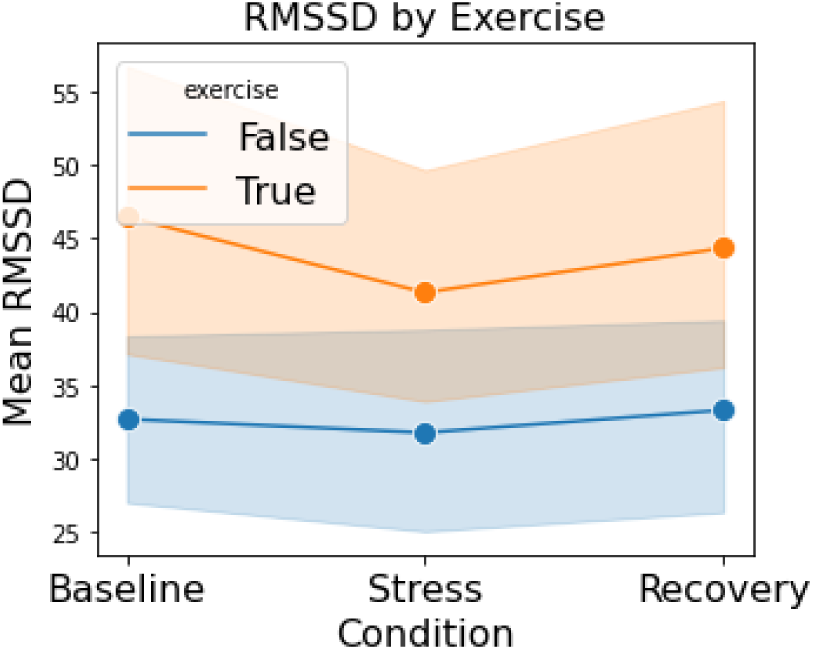
RMSSD Across Experimental Conditions, divided by exercise regularly (‘True’ for at least 1-2 times a week vs. ‘False’ for none).

#### Individual Characteristics Modulating Self-Reported Stress

A significant effect of sex was identified, with males self-reporting significantly lower stress levels compared to females (β −1.200, p = 0.028). See Figure 10.

**Figure 10.**
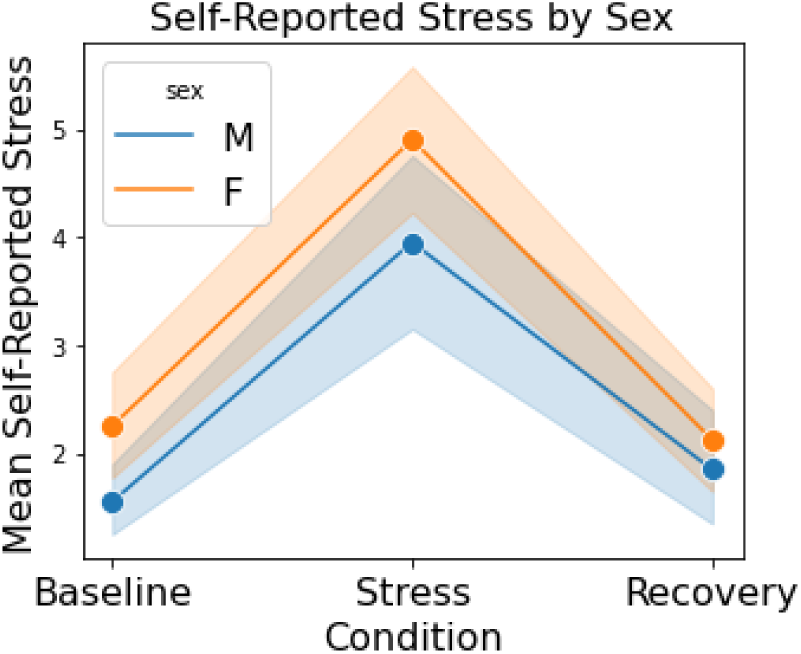
Self-Reported Stress Across Experimental Conditions, divided by Sex (Female vs. Male).

## Discussion

The present study aimed to assess the stress level scores during varying stress/rest conditions, provided by the Garmin Vivosmart 4 fitness tracker. GSS values were tested against HR and HRV parameters obtained, from what is considered a gold standard, Polar H10 chest strap recordings. Participants were exposed to three conditions: baseline (rest), stress (arithmetic task), and recovery (relaxing video/music). Each of the experimental conditions lasted 15 minutes during the main experiment, and 30 minutes during an earlier pilot study. The primary objective of the analysis was to identify and evaluate physiological responses across conditions encompassing variations in both HR and HRV metrics, in addition to GSSs, alongside subjective self-reported stress. We measured the strength and direction of correlations between the stress score given by the Garmin device and HR and HRV metrics.

Our pre-registered analysis (based on the pilot study) revealed a dynamic response of physiological parameters to the stress-inducing task, as evidenced by significant differences in mean HR, SD2/SD1 ratio, and HF power (nu) between stress conditions and baseline/rest periods. In addition, subjective self-reported stress levels were higher at the end of the stress-inducing task, supporting protocol efficacy. These results align with previous studies indicating alterations in autonomic balance during mental stress [12], specifically increased sympathetic activity and decreased parasympathetic activity. GSSs were significantly higher during the stress condition relative to baseline and recovery phases, demonstrating that it successfully reflects the stress response. Additionally, our data revealed associations between GSS and the physiological parameters, specifically a high correlation with HR and moderate correlations with SD2/SD1 and RMSSD. These results shed light on the potential utility of the wearable device in assessing stress levels remotely in natural settings.

### Physiological Responses to Mental Stress Task and GSS Trends

Both the pilot and pre-registered (main) experiment consistently observed a parallel pattern between GSS and mean HR across the baseline, stress, and recovery phases, demonstrating a significant increase during the stress task and a decrease during recovery. Furthermore, among HRV metrics that were assessed (RMSSD, SD2/SD1, LF/HF, HF power), SD2/SD1 exhibited a significant increase during the stress task in both the pilot and main experiments. However, changes in RMSSD were not statistically significant. Among the spectral analysis metrics, results from the main experiment demonstrated a significant decrease in HF Power during the stress condition, while changes in LF/HF ratios were not statistically significant in either experiment. Furthermore, self-reported subjective stress levels reported by participants after the stress-inducing task were significantly higher compared to levels reported right after both baseline and recovery phases, supporting the efficacy of the arithmetic task as an effective tool for eliciting perceived stress among participants.

The significant increase in mean HR during the stress task compared to baseline and recovery is consistent with sympathetic nervous system activation commonly associated with heightened physiological arousal or stress responses [12], [65], [66]. The decrease in HF power during the stress condition is consistent with parasympathetic nervous system withdrawal associated with stress response [12], [18], [29], [67].

An increase in SD2, associated with long-term variability, may indicate sympathetic activation, while decreases in SD1, linked to short-term variability, may reflect reduced parasympathetic activity [68]. Therefore, an elevated SD2/SD1 ratio could suggest a shift in autonomic balance towards sympathetic dominance.

The lack of significant difference in RMSSD and LF/HF across conditions in the univariate analysis, in both the pilot and pre-registered experiments, warrants further investigation into the sensitivity of these metrics to the chosen stress task or to the effect of individual differences. RMSSD, as a measure of HRV, reflects the beat-to-beat variation in heart rate and is predominantly influenced by parasympathetic nervous system activity [18]. The lack of significant findings regarding RMSSD could be attributed to various factors including stress response complexity and individual differences. Although stress typically correlates with reduced parasympathetic activity and subsequent RMSSD decreases, the stress response is multifaceted and may involve intricate physiological mechanisms. For instance, one study [69] illustrated an interesting interplay between tonic HRV and phasic HRV and demonstrated how tonic HRV might affect the suppression or enhancement of phasic HRV. This dynamic could also extend to how individual differences in stress reactivity and stress regulation (as well as coping mechanisms) might influence RMSSD levels during stressful tasks, leading to variability in responses within the study sample. Additionally, temporal dynamics could offer an alternative explanation. RMSSD reflects short-term variations in heart rate, influenced by rapid changes in autonomic activity [23]. Stress-induced changes in autonomic balance may have been transient or occurred at different time points during the stress task, possibly contributing to the lack of significant differences in RMSSD between conditions. Despite the absence of statistical significance, the trend of lower RMSSD during the stress condition compared to baseline aligns with theoretical expectations and previous research on stress-induced HRV changes [12]. While the lack of statistical significance may limit the interpretability of these findings, they nonetheless contribute to our understanding of the complex interplay between stress and autonomic regulation.

The complexity of stress response and individual differences may contribute also to the inconsistent differences observed in LF/HF across conditions. Moreover, critics have challenged the view that the LF/HF ratio reflects a sympatho-vagal balance due to its weak correlation with sympathetic nerve activation and the non-linear relationship between sympathetic and parasympathetic activity [27], [28]. We illustrated the impact of sex on LF/HF and HF metrics. The observation that these metrics change in opposite directions in response to stressors could potentially explain the mixed results. Previous research that investigated HRV changes during mental stress has similarly reported non-significant differences in LF/HF ratio [65]. Therefore, although this metric is frequently utilized in HRV research, its reliability remains uncertain and as such, its impact on the interpretation of our findings.

### Correlations of Garmin metric with HRV and mean HR

The correlational analysis revealed a consistently high positive correlation between the GSS and mean HR as evident in both computation methods (average of within-subject correlations and within-condition (between-subject analysis). Results were consistent across both the pilot and the pre-registered experiment. Conversely, correlations between the GSS and HRV metrics were comparatively lower, yet significant. In the main experiment, the Garmin metric exhibited a negative correlation with RMSSD and a positive correlation with SD2/SD1, aligning with the expected autonomic shift in response to the stress task. Conversely, associations with spectral analysis metrics (LF/HF and HF power) remained low across both experiments and computation methods, mirroring the lack of significant change observed in the variations of measures across conditions analysis.

The strong correlation of the GSS with HR might suggests that HR may carry greater weight than HRV in the Garmin algorithm’s stress calculation. The high standard deviations in the correlation analyses suggest considerable variability among participants. This variability may reflect individual differences in autonomic regulation and stress responses, highlighting the need for personalized approaches in HRV analysis.

### Outliers Exclusion

The exclusion of outliers resulted in minor changes to descriptive statistics that did not alter the interpretation of our findings. This suggests that the underlying trends and relationships in the dataset were robust to the presence of extreme observations. The cleaned dataset continued to demonstrate the same patterns as the original data, affirming the reliability of the observed effects.

### Exploratory Analysis: The Role of Individual Differences in Modulating GSS, HR, And HRV

We further explored the effects of individual characteristics, including sex, age, BMI, habitual exercise, average nightly sleep duration, and tonic HRV (low vs. high RMSSD levels at baseline) on GSS, HR, HRV measures, and self-reported stress. Using mixed-effect linear model analysis we demonstrated how sex, exercise, and tonic HRV significantly modulate HR and HRV values. while GSS was influenced by exercise, tonic HRV, BMI, and average nightly sleep duration.

### The Effect of Tonic HRV

Existing research demonstrates that individuals with higher tonic HRV exhibit more adaptive stress responses than those with lower HRV [12], [15], as higher HRV reflects greater parasympathetic (vagal) tone, which facilitates effective stress regulation, while lower HRV indicates reduced vagal influence and impaired stress adaptation [70]. This aligns with our findings, where participants with higher baseline HRV displayed significantly lower HR and lower GSS overall, especially during the stress task. An interaction between baseline HRV and the stress condition reveals opposite trends across conditions: high HRV individuals show a decrease in RMSSD during stress and a rebound during recovery, while low HRV individuals do not exhibit this pattern. However, while a significant difference between groups was observed, the changes between conditions in the high HRV group were not statistically significant, limiting the conclusions that can be drawn from these results.

### The Effect of Sex

Sex differences were exhibited specifically in the frequency domain and self-reported stress metrics. While males self-reported lower stress, they displayed higher LF/HF values and lower HF values compared to females. These findings align with past data supporting sex differences in HRV, with women having greater HRV than men even after controlling for many potential confounders. It was demonstrated in a twin study which investigated the effect of sex differences and heritability on heart rate dynamics [71].

The results indicate that the impact of stress on the SD2/SD1 ratio and HFnu differs between males and females, as evidenced by a significant interaction effect between sex and the stress condition. Analysis of frequency domain metrics revealed that in reaction to stress, women exhibit an increase in LF/HF and a decrease in HFnu, while men exhibited an opposite pattern. The trend displayed by women in our experiment was consistent with existing literature involving the effect of stress on HRV [12]. These findings contribute to the understanding of the results that emerged from the pre-registered analysis. Using a univariate analysis (paired t-test), without controlling for individual characteristics, the comparison of the frequency domain HRV metrics across conditions displayed a relatively small effect to non-significant. The disparity between sexes in how LF/HF alters in response to stress might have contributed to the insignificant differences it exhibited in the univariate analysis which did not control for sex. This may also explain the relative weak correlations of LF/HF and HFnu with the GSS that our data displayed. Given that Garmin gathers data on factors like sex, age, fitness level, and sleep, we presume it incorporates them into its stress computation.

In a meta-analysis by Koenig and Thayer [33] on sex differences in HRV, it was found that females typically have a higher average heart rate, as indicated by a shorter mean RR interval. In the frequency domain, females exhibited greater power in the high-frequency (HF) band. These results suggest that the autonomic regulation of the female heart is predominantly governed by vagal, parasympathetic activity, even with a higher average heart rate. In contrast, the male heart shows relative sympathetic dominance, despite having a lower heart rate.

### The Effect of Exercise

Our findings also suggest that individuals who exercise at least 1-2 times a week have, on average, a lower HR and a lower GSS compared to those who do not exercise regularly. Those findings are aligned with the existing literature, associating higher HRV and lower HR with higher fitness levels [34]. Regarding whether habitual exercise influences the response to mental stress as reflected through HR and HRV, we did not observe a significant effect. In both groups, those who exercise regularly and those who do not, the physiological variables demonstrated a parallel pattern of the stress response, including the GSS.

### Stress Task Effect

The LMM identified a significant fixed effect of the stress condition on GSS, self-reported stress, HR, and HRV metrics (excluding LF/HF), while controlling for individual characteristics. Additionally, we identified higher levels of GSS during the **recovery condition**, compared to baseline, indicating residual stress effects post-task. A similar post-stress-task effect was observed in HFnu values, with a decreased level in the recovery phase compared to baseline. These findings add and strengthen the results of the pre-registered univariate analysis (paired t-test) and confirm the stress-induced task’s efficacy. Additionally, they demonstrate the GSS’s capability to detect stress levels comparable to HR and certain HRV measurements.

While our findings provide valuable insights, several limitations must be acknowledged, and further exploration could provide additional evidence concerning the effectiveness of a stress score based on HRV and the GSS metrics in assessing individuals’ stress responses:

#### The Stress Induction Method

We relied on an arithmetic quiz to induce mental stress. Future studies could explore different stress-induction techniques for a more comprehensive understanding.

#### Longitudinal Studies

Longitudinal studies tracking GSS and HRV patterns over time in response to chronic stressors could provide valuable insights into the long-term impact of stress on autonomic function and assess the effectiveness of the GSS in monitoring stress variations over time. Van Kraaij and colleagues [72] found a significant relationship between chronic stress and heart rate over time using various wearable devices. Limitations of the current study are the use of a single-event intervention to induce stress, which may not have activated stress sufficiently in all participants. Future research should consider employing a longitudinal experimental design to examine the impact of accumulating stressors over time. Such an approach would allow for a more comprehensive understanding of stress responses and their physiological manifestations.

#### Additional stress measures

Future studies could incorporate additional stress measures (e.g., cortisol levels) to be compared with the wearable metric and provide a wider picture of the physiological response to the stressor.

#### Moderating variables

Future research should further investigate moderating variables in stress reactivity to deepen understanding of how individual differences shape stress responses, as these factors appear to play a significant role in modulating both the experience and physiological impact of stress on individuals. These could include factors such as age, sex, personality traits (e.g., neuroticism or resilience), baseline fitness levels, chronic stress exposure, sleep quality, and lifestyle behaviors (e.g., caffeine consumption or physical activity patterns).

#### Chronic Disease Monitoring

Given the association between the stress level score and HRV metrics in the current study, alongside the established evidence of low vagally-mediated HRV as a marker of health risks [21], [22], it is worth exploring the potential use of Garmin device for monitoring patients with existing health conditions. Continuous, real-time tracking of psychobiological changes using the Garmin stress score could provide insights into the dynamics of disease progression, including stress responses and comorbidities. Such information may help to better understand underlying physiological processes and serve as an early-warning system to prevent exacerbation of chronic diseases through timely, tailored interventions.

## Conclusion

Our findings underscore the dynamic nature of autonomic nervous system regulation in response to mental stressors, revealing significant variations in physiological responses. The significant elevation in GSS during the stressful task, coupled with its high association with HR and moderate correlations with certain HRV metrics, calculated by Kubios HRV based on measurements with a Polar H10 chest strap, suggests that the device effectively detects the physiological response to the mental stress task used in this study.

Elevated HR and sympathetic activity during stress, alongside reduced parasympathetic activity indicated by decreased HF power, were observed. GSS mirrored these patterns. Specifically, GSS exhibited a significant increase during stress compared to baseline. Additionally, tonic HRV, exercise, BMI, and sleep duration significantly influenced GSS, with high tonic HRV, regular exercise, and longer sleep duration associated with lower stress scores, while higher BMI and low tonic HRV were linked to higher stress scores. The observed correlations between the GSS and HR and HRV metrics, alongside consistent findings between the pilot and main pre-registered experiment, support the Garmin Vivosmart 4 as a potential tool for mental stress level monitoring. The observed associations highlight the potential of wearable technology, in objectively assessing stress levels in real-world settings. Their noninvasive and remote nature facilitates longitudinal and ecological research in health, behavior, and other fields of human research. Accessibility and ease of use enable large-scale studies, facilitating exploration across diverse populations and contexts. Furthermore, integrating GSS with other physiological and behavioral measurements could provide a comprehensive understanding of stress-related processes and their impact on behavior, health, and well-being.

Future research encompassing a broader range of stress induction methods, and longitudinal designs with consideration of individual differences, could further enhance our understanding of the GSS’s utility in real-world stress management, mental and physical alike.

## Supporting information

Supplemental Section A

Supplemental Section B

Supplemental Section C Table S IV

Supplemental Section C Table S V

Supplemental Section C Table S VI

Supplemental Section C Table S VII

Supplemental Section C Table S VIII

Supplemental Section C Table S IX

## Acknowledgment

We would like to thank Dana Roll for managing the experiment’s execution and research team, and overseeing participant recruitment, as well as Danielle David, Li-Or Oren, and Liel Cohen for their invaluable assistance in running the experimental procedure and recruiting participants. We are also grateful to the participants who agreed to be part of this study. Additionally, we extend our gratitude to Jeanette Mumford for her expert advice in statistical analysis.

## Notes

### Competing Interest Statement

The authors have declared no competing interest.

