## Supplemental Section A for "Assessing Garmin’s Stress Level Score Against Heart Rate Variability Measurements"

### Section A: Pilot Experiment - Methods

#### Pilot Experiment - Participants

Participant recruitment and inclusion: a total of 35 participants (24 females) with a mean age of 26.4 years (SD = 5.04 years; range: 21-39) volunteered for the experiment. Participants were University students or those who responded to social media advertisements. All participants were healthy, had normal or corrected-to-normal vision, and normal hearing, reported no neurological conditions, did not consume any psychiatric medications or drugs, and were not diagnosed as suffering from cardiovascular irregularities. Women declared that they were not pregnant. Data from six participants was excluded from the analysis due to device failures during the procedure, resulting in a final valid sample size of 29. Ethical approval was obtained from the ethics committee of Tel Aviv University, and all participants provided informed consent before participation. Participants were compensated by a payment of 40 NIS per hour. In addition, a bonus payment ranging between 0-20 NIS was awarded according to individual success in the stress task.

#### Pilot Experiment - Procedure

To explore the relationship between HRV parameters and stress level scores obtained from the Garmin Vivosmart 4 (GV4) fitness tracker, data were collected concurrently using the GV4 and a Polar H10 chest-strap heart rate monitor (Polar Electro Oy) during the main laboratory session. The Polar H10 served as the validation device for recording RR signals extracted from the QRS complex using electrocardiography (ECG) technology. Stress level scores were calculated by the GV4 which uses a photoplethysmography (PPG) sensor.

The procedure included two laboratory sessions for each participant, a preparatory meeting, and an experimental task session. During a preparatory meeting, participants were provided with a GV4 fitness tracker, which was synchronized with a designated personal account in the Garmin Connect mobile application on their phones. Additionally, demographic information including age, sex, height, and weight were collected, and inclusion criteria were verified. Informed consent was obtained after explaining the experimental procedures. To ensure accurate identification of stressful moments, participants were instructed to wear the fitness tracker for one full day and night prior to the main laboratory session, as per the recommendations outlined in the GV4 manual. Participants were instructed to abstain from food, caffeine, and smoking for at least two hours before the experimental task session, and to refrain from intense physical activity 24 hours prior to the session.

Participants came back to the laboratory for the experimental task session the following day or later in the week. At the beginning of the session, the experimenter ensured that stress and heart rate data had been collected using the GV4 during the preceding day and night. This session comprised three consecutive tasks, each lasting 30 minutes, conducted in the laboratory: a resting state (baseline), a stressful state, and a relaxation state (recovery), amounting to a total duration of 90 minutes. Throughout the session, participants remained seated in front of a computer in a quiet laboratory testing room, wearing both the GV4 fitness tracker and the Polar H10 chest strap. They were instructed to minimize movement during the tasks while data were simultaneously collected from both devices. Before the task, participants were informed of a monetary incentive contingent upon their performance. During the resting and relaxation states, participants viewed a video featuring natural landscapes and listened to calming music. In the stressful state, participants engaged in a computerized mental arithmetic task modeled after the "Montreal Imaging Stress Task" (MIST), a protocol used to induce psychosocial stress in participants [53]. Following the completion of the three tasks, participants completed a Perceived Stress Scale questionnaire (PSS-14), [54] after the entire 90-minute data collection. Figure  provides a visual representation of the experimental protocol.

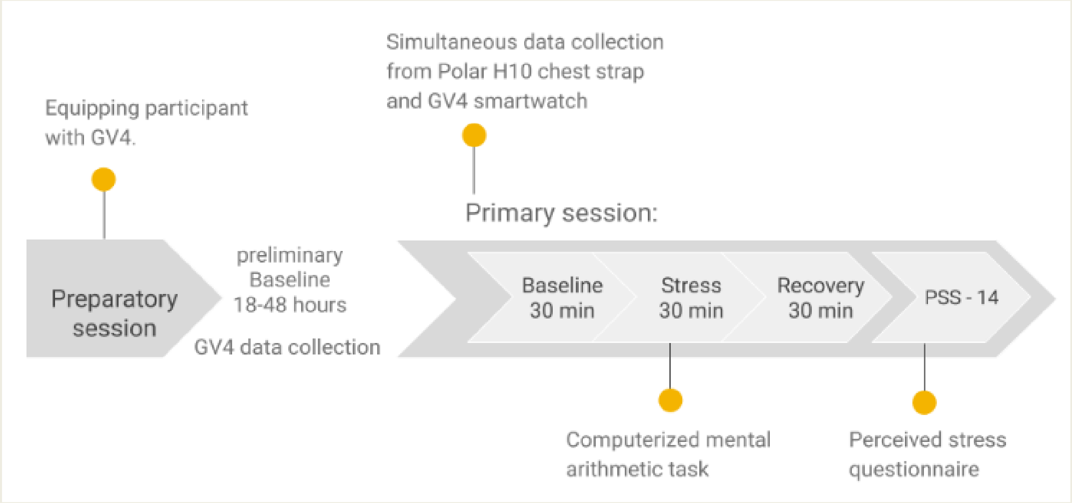


Figure S1. Protocol Flowchart of the Pilot Study.

#### Pilot Experiment - Data Analysis

The dataset obtained in the pilot consisted of twenty-nine participants, with each participant providing thirty 3-minute interval observations. Each observation included mean HR, HRV metrics (e.g., RMSSD [ms], HF power [nu]) derived from an IBI signal recorded by the Polar H10 chest strap, and stress scores from the Garmin fitness tracker. This resulted in a total of 90 minutes of data per participant, with each variable represented by a column in the dataset table. The analysis involved assessing the mean differences between stress conditions and both baseline and recovery phases, as well as computing the correlation between GSSs and selected HRV metrics. The selected metrics evaluated with GSS included mean HR (bpm), RMSSD (ms), SD2/SD1, HF power (nu), and LF/HF. Statistical significance was determined with p values $< 0.05$. To account for multiple comparisons (of HRV metrics), the Bonferroni correction was applied, setting the significance level to $p < 0.05/4 = 0.0125$.

We first identified stress-related physiological responses during the stress condition by comparing Mean values of HRV parameters, HR, and GSSs across baseline, stress, and recovery conditions. Mean values were calculated for every metric in each condition, and paired t-tests were utilized to compare these values across the different conditions. To examine the relationship between GSSs and HRV metrics, we employed two methods of correlation analysis. The first method involved a correlation analysis of condition means. For this analysis, we calculated the mean GSSs and HRV metrics within each experimental condition (baseline, stress, and recovery) for each participant. Using these means, we computed the Pearson correlation coefficient for each condition based on the data from all 29 participants. This method allowed us to assess the strength and direction of the relationship between HR, HRV parameters, and GSSs across different states. The second method was a within-subject analysis. For each participant, we calculated the Pearson correlation coefficient between GSSs and each HRV metric. We then averaged these individual correlation coefficients across all participants to obtain a mean correlation coefficient for each HRV metric. To determine the significance of these correlations, we conducted a one-sample t-test on the mean correlation coefficients. This process resulted in a table displaying the mean Pearson's r and associated p-values for each HRV metric with GSSs, indicating whether there was a meaningful relationship between GSSs and HRV metrics across the participant sample. This two-pronged approach provided a comprehensive evaluation of the association between GSSs and HRV metrics, both at the group level and within individual participants.
