## Supplemental Section B for "Assessing Garmin’s Stress Level Score Against Heart Rate Variability Measurements"

### Section B: Pilot Experiment – Results

#### Pilot Experiment Results – Participants

Data from 29 participants (19 females), with a mean age of 25.8 years (SD = 4.2), were valid for analysis. Demographic and other information is provided in Table S I.

Table S I: Participant information - Pilot Study

| | Mean | Std | Min | Max | Count | Percentage |
|---|---|---|---|---|---|---|
| Age [years] | 25.8 | 4.2 | 21 | 38 | - | - |
| Height [cm] | 167.4 | 7.9 | 155 | 184 | - | - |
| Weight [Kg] | 62.1 | 11.1 | 42 | 90 | - | - |
| Sex | | | | | | |
| Female | - | - | | | 19 | 65.5% |
| Male | - | - | | | 10 | 34.5% |
| PSS-14 | 24.6 | 8.5 | 4 | 43 | - | - |

#### Pilot Experiment Results – Variation of Measures Across Experimental Conditions

*Stress Task Effects*: paired t-tests were conducted to assess significant differences in participants' physiological measures across the three conditions (Baseline, Stress, and Recovery). A significance threshold was set at $p < 0.0125$ after applying the Bonferroni correction to account for multiple comparisons of HRV metrics.

The results of this analysis revealed significant differences in Mean HR between Stress and Baseline ($p < 0.001$) and between Stress and Recovery ($p < 0.001$), indicating elevated HR during the stress-inducing task compared to rest periods. No significant differences were observed in the comparison of HR during Recovery vs HR during Baseline. Mean RMSSD increased during the recovery condition relative to the stress and baseline, however, this difference did not reach the significant threshold. The SD2/SD1 Ratio exhibits significant difference between Stress and Baseline ($p = 0.0045$). LF/HF Ratio and HF Power (nu) did not exhibit significant differences between the three conditions, although HF power did exhibit a decrease during the stress condition relative to baseline with a minor increase during recovery. The GSS showed a very similar pattern to the HR with a significant increase during the stress task ($p < 0.001$) and a decrease during recovery ($p < 0.001$). These findings underscore the dynamic physiological responses to stress-inducing tasks and subsequent recovery periods, see Figure S2 for a visual presentation of the comparison. For full results, please refer to

Table S II.

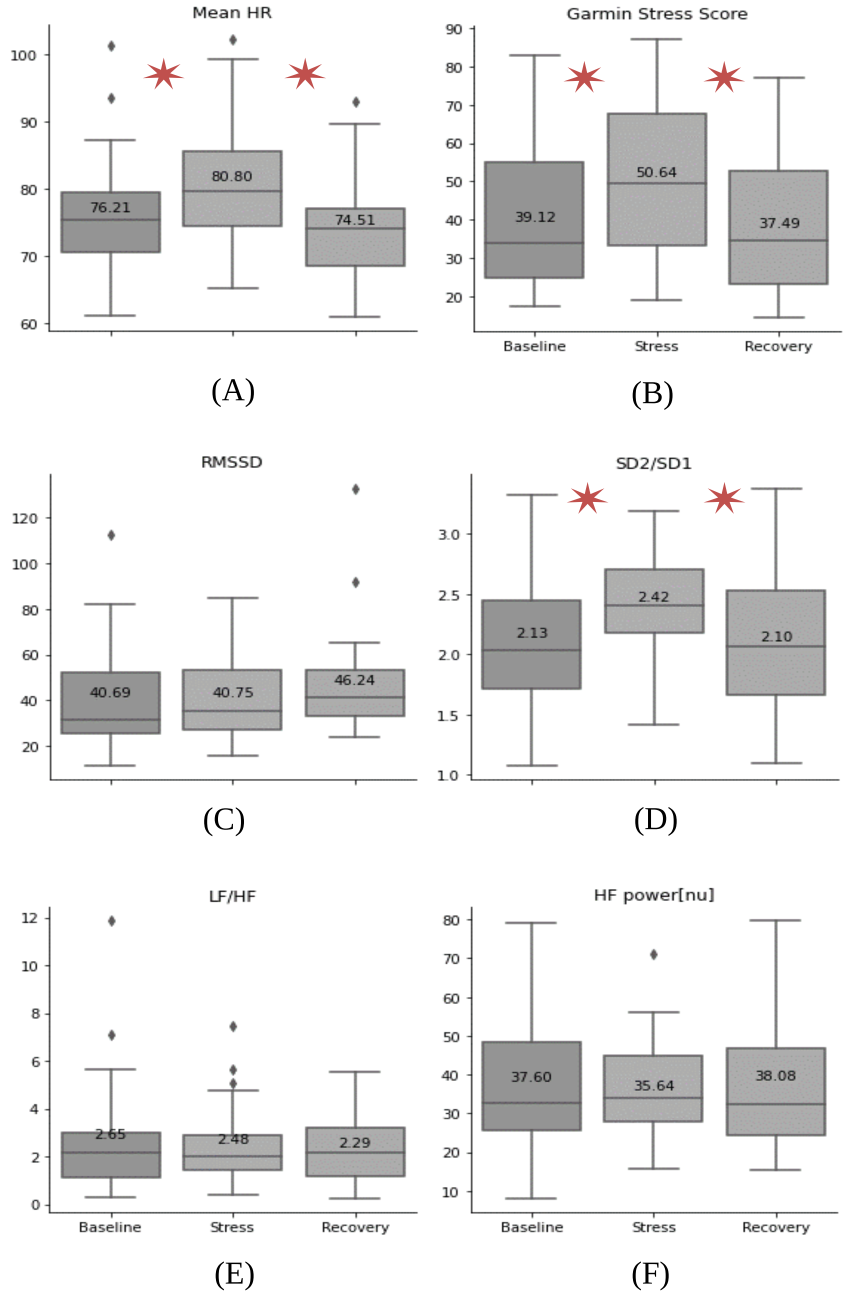


Figure S2. Distribution of HR, HRV, and GSSs across the baseline (rest), stress, and recovery conditions in the pilot. Y axes represent the value of each parameter. Notably, mean values of each condition are indicated in the boxplot. Significant differences (P value < 0.0125) across conditions are marked by a star.

Table S II: Descriptive Statistics of Metrics Under the Three Experimental Conditions and Paired T-Test Results of Comparisons - Pilot

| Condition | BASELINE | | STRESS | | RECOVERY | | STRESS VS. BASELINE | | RECOVERY VS. BASELINE | | RECOVERY VS. STRESS | |
|---|---|---|---|---|---|---|---|---|---|---|---|---|
| | Mean | SD | Mean | SD | Mean | SD | Delta | p-value | Delta | p-value | Delta | p-value |
| **Mean HR (bpm)** | 76.21 | (8.73) | 80.8 | (8.84) | 74.51 | (7.93) | 4.59 ** | 2.53E-05 | -1.7 | 0.040 | -6.29 ** | 5.20E-06 |
| **RMSSD (ms)** | 40.69 | (22.01) | 40.75 | (18.57) | 46.24 | (22.06) | 0.06 | 0.976 | 5.55 * | 0.007 | 5.49 | 0.028 |
| **SD2/SD1** | 2.13 | (0.61) | 2.42 | (0.5) | 2.1 | (0.57) | 0.29 * | 0.005 | -0.03 | 0.725 | -0.32 * | 0.001 |
| **LF/HF** | 2.65 | (2.41) | 2.48 | (1.63) | 2.29 | (1.47) | -0.17 | 0.679 | -0.36 | 0.291 | -0.19 | 0.432 |
| **HF (nu)** | 37.6 | (18.7) | 35.64 | (13.47) | 38.08 | (17.08) | -1.96 | 0.465 | 0.48 | 0.799 | 2.44 | 0.293 |
| **GSS** | 39.12 | (17.43) | 50.64 | (19.64) | 37.49 | (18.33) | 11.52 ** | 8.60E-07 | -1.63 | 0.507 | -13.15 ** | 7.67E-06 |

* Note: * P < 0.0125, ** P < 0.001

#### Pilot Experiment Results – Correlation Analysis

Correlation analysis results revealed consistent associations between GSSs and various physiological metrics, employing two different methods of computation.

Within-Condition Correlation (analysis of condition means)

In Method 1, Pearson correlation coefficients were computed from mean scores across participants, within each condition (each participant had one observation in each condition, totaling 3 observations for participant). Strong positive correlations emerged between GSSs and Mean HR in three conditions, with coefficients ranging from 0.69 to 0.75. Slightly lower positive correlation coefficients were observed between Garmin's metric to SD2/SD1 (0.48 to 0.52). Conversely, a moderate negative correlation ($r = -0.47$, $p < 0.01$) between RMSSD and Garmin's metric was significant for the baseline measurement only. Furthermore, correlations between GSSs and LF/HF were found to be lower, ranging from 0.2 to 0.35. HF power exhibited a significant correlation with Garmin's metric for the baseline measurement ($r = -0.48$, $p < 0.01$). See Table S III for Pearson Correlation Coefficients in each condition and corresponding P-values.

Table S III: Pearson Correlation Coefficient between GSS and HRV Metrics and mean HR in each Experimental Condition - Pilot

| CONDITION | BASELINE | | | STRESS | | | RECOVERY | | |
|---|---|---|---|---|---|---|---|---|---|
| | Pearson r | p-value | sig | Pearson r | p-value | sig | Pearson r | p-value | sig |
| **MEAN HR (bpm)** | 0.75 | 3.08E-06 | * | 0.76 | 1.89E-06 | * | 0.69 | 3.94E-05 | * |
| **RMSSD (ms)** | -0.47 | 0.00935 | * | -0.46 | 0.012714 | | -0.4 | 0.032293 | |
| **SD2/SD1** | 0.48 | 0.009109 | * | 0.48 | 0.008022 | * | 0.52 | 0.003972 | * |
| **LF/HF** | 0.35 | 0.060313 | | 0.2 | 0.290939 | | 0.35 | 0.064578 | |
| **HF power (nu)** | -0.48 | 0.0089 | * | -0.37 | 0.048895 | | -0.45 | 0.014713 | |

Note: significant results (p value < 0.0125) are marked by a star (*).

##### *Within-Subject Correlation Analysis*

In Method 2, we investigated the relationships between GSSs and physiological parameters at the individual level. Therefore, we computed individual correlations for each participant and analyzed the averages across all participants. This method revealed significant mean correlation coefficients across all metrics tested with GSSs, except for HF power. Specifically, Mean HR exhibited a relatively strong positive correlation ($r = 0.72$, $SD = 0.29$, $p < 0.001$), indicating a consistent link between higher stress scores and elevated heart rates. RMSSD displayed a moderate negative correlation ($r = -0.33$, $SD = 0.42$, $p < 0.001$), suggesting that increased stress scores were associated with reduced heart rate variability. SD2/SD1 demonstrated a moderate positive correlation ($r = 0.38$, $SD = 0.28$, $p < 0.001$), indicating a tendency for higher stress scores to align with higher SD2/SD1 ratios. In contrast, LF/HF and HF exhibited low correlations with Pearson’s r of 0.20 ($SD = 0.33$, $p = 0.004$) and -0.18 ($SD = 0.37$, $p = 0.013$), respectively, indicating weaker associations with Garmin's metric. High standard deviations were observed in the correlation statistics, indicating variability in these measures among participants.
