## Supplemental Section C Table S IV for "Assessing Garmin’s Stress Level Score Against Heart Rate Variability Measurements"

### Section C: Main Experiment – Additional Results

Table S I: Descriptive statistics of Metrics Under the Three Experimental Conditions and paired t-test results of comparisons - Main Experiment

| CONDITION | BASELINE | | STRESS | | RECOVERY | | STRESS - BASELINE | | RECOVERY - BASELINE | | RECOVERY - STRESS | |
|---|---|---|---|---|---|---|---|---|---|---|---|---|
| | Mean | SD | Mean | SD | Mean | SD | Delta | p-value | Delta | p-value | Delta | p-value |
| **MEAN HR (bpm)** | 73.93 | (12.55) | 78.93 | (12.7) | 74.29 | (12.01) | 5 ** | 1.43E-09 | 0.36 | 0.430 | -4.64 ** | 1.73E-11 |
| **RMSSD (ms)** | 41.2 | (25.91) | 37.66 | (22.81) | 40.12 | (24.92) | -3.54 | 0.054 | -1.08 | 0.429 | 2.46 | 0.047 |
| **SD2/SD1** | 2.19 | (0.71) | 2.41 | (0.67) | 2.3 | (0.76) | 0.22 * | 0.001 | 0.11 | 0.014 | -0.11 | 0.069 |
| **LF/HF** | 2.63 | (2.3) | 2.91 | (2.3) | 2.91 | (2.56) | 0.28 | 0.210 | 0.28 | 0.172 | 0 | 0.975 |
| **HF (nu)** | 38.61 | (17.08) | 33.54 | (13.91) | 35.76 | (15.88) | -5.07 * | 0.003 | -2.85 | 0.035 | 2.22 | 0.087 |
| **GSS** | 41.9 | (24.31) | 50.16 | (25.54) | 43.63 | (24.33) | 8.26 ** | 7.03E-08 | 1.73 | 0.182 | -6.53 ** | 1.88E-07 |
| **SELF-REPORTED STRESS** | 2.02 | (1.36) | 4.58 | (2.13) | 2.03 | (1.45) | 2.56 * | 1.51E-13 | 0.01 | 0.929 | -2.55 * | 1.32E-11 |

NOTE: *P < 0.0125, ** P < 0.001
