## Supplemental Section C Table S V for "Assessing Garmin’s Stress Level Score Against Heart Rate Variability Measurements"

### Section C: Main Experiment – Additional Results

Table S I: Pearson Correlation Coefficient between HR and HRV Metrics and GSS in each Experimental Condition - Main Experiment

| **CONDITION** | **BASELINE** | | | **STRESS** | | | **RECOVERY** | | |
|---|---|---|---|---|---|---|---|---|---|
| | Pearson r | p-value | sig | Pearson r | p-value | sig | Pearson r | p-value | sig |
| **MEAN HR (bpm)** | 0.85 | 1.01E-17 | * | 0.84 | 5.53E-17 | * | 0.85 | 1.56E-17 | * |
| **RMSSD (ms)** | -0.6 | 3.71E-07 | * | -0.63 | 5.08E-08 | * | -0.59 | 6.18E-07 | * |
| **SD2/SD1** | 0.62 | 1.20E-07 | * | 0.64 | 2.73E-08 | * | 0.61 | 2.16E-07 | * |
| **LF/HF** | 0.38 | 0.0028 | * | 0.32 | 0.0126 | | 0.35 | 0.0065 | * |
| **HF (nu)** | -0.4 | 0.0014 | * | -0.43 | 0.0006 | * | -0.43 | 0.0007 | * |

* Note: *P < 0.0125
