## Supplemental Section C Table S VI for "Assessing Garmin’s Stress Level Score Against Heart Rate Variability Measurements"

### Section C: Main Experiment – Additional Results

Table S I: Pearson Correlation Coefficient between GSS and HR/HRV Metrics - Within-subject Analysis

| **Mean Pearson r: GSS and HR,HRV** | |
|---|---|
| Mean HR (bpm) | 0.74 * |
| RMSSD (ms) | -0.41 * |
| SD2/SD1 Ratio | 0.3 * |
| LF/HF Ratio | 0.18 * |
| HF Power (nu) | -0.2 * |

* Note: *P < 0.0125
