## Supplemental Section C Table S VII for "Assessing Garmin’s Stress Level Score Against Heart Rate Variability Measurements"

### Section C: Main Experiment – Additional Results

#### Analyses Post-Outlier Removal

Table S I: Descriptive statistics of Metrics Under the Three Experimental Conditions and paired t-test results of comparisons – Cleaned Dataset (after the removal of outliers)

| **CONDITION** | **BASELINE** | | **STRESS** | | **RECOVERY** | | **STRESS - BASELINE** | | **RECOVERY - BASELINE** | | **RECOVERY - STRESS** | |
|---|---|---|---|---|---|---|---|---|---|---|---|---|
| | Mean | SD | Mean | SD | Mean | SD | Delta | p-value | Delta | p-value | Delta | p-value |
| **MEAN HR (bpm)** | 73.36 | (11.8) | 78.21 | (11.26) | 73.78 | (11.2) | 4.85** | 3.03E-09 | 0.42 | 0.363 | -4.43** | 2.95E-11 |
| **RMSSD (ms)** | 41.4 | (24.77) | 37.41 | (19.56) | 39.44 | (21.18) | -3.99 | 0.033 | -1.96 | 0.115 | 2.03 | 0.072 |
| **SD2/SD1** | 2.15 | (0.65) | 2.37 | (0.61) | 2.27 | (0.71) | 0.22* | 0.001 | 0.12* | 0.008 | -0.1 | 0.087 |
| **LF/HF** | 2.39 | (1.9) | 2.71 | (1.82) | 2.77 | (2.25) | 0.32 | 0.106 | 0.38 | 0.028 | 0.06 | 0.741 |
| **HF (nu)** | 39.02 | (16.19) | 33.68 | (12.92) | 36.23 | (15.96) | -5.34* | 0.002 | -2.79 | 0.045 | 2.55 | 0.062 |
| **GSS** | 41.09 | (23.51) | 49.43 | (24.85) | 42.74 | (23.63) | 8.34** | 8.32E-08 | 1.65 | 0.211 | -6.69** | 1.43E-07 |
| **SELF-REPORTED STRESS** | 1.97 | (1.31) | 4.58 | (2.15) | 1.98 | (1.41) | 2.61** | 1.12E-13 | 0.01 | 0.929 | -2.6** | 1.08E-11 |

* Note: *P < 0.0125
