## Supplemental Section C Table S VIII for "Assessing Garmin’s Stress Level Score Against Heart Rate Variability Measurements"

### Section C: Main Experiment – Additional Results

#### Analyses Post-Outlier Removal

Table S I: Pearson Correlation Coefficient between HR and HRV Metrics and GSS in each Experimental Condition – Cleaned Dataset (after the removal of outliers)

| **CONDITION** | **BASELINE** | | | **STRESS** | | | **RECOVERY** | | |
|---|---|---|---|---|---|---|---|---|---|
| | Pearson r | p-value | sig | Pearson r | p-value | sig | Pearson r | p-value | sig |
| **MEAN HR (bpm)** | 0.84 | 1.94E-16 | * | 0.83 | 6.07E-16 | * | 0.84 | 1.24E-16 | 0.84 |
| **RMSSD (ms)** | -0.59 | 8.01E-07 | * | -0.58 | 1.21E-06 | * | -0.63 | 7.68E-08 | -0.59 |
| **SD2/SD1** | 0.59 | 7.46E-07 | * | 0.57 | 2.09E-06 | * | 0.62 | 1.84E-07 | 0.59 |
| **LF/HF** | 0.33 | 0.012 | * | 0.3 | 0.019 | | 0.31 | 0.016 | 0.33 |
| **HF (nu)** | -0.35 | 0.006 | * | -0.4 | 0.002 | * | -0.38 | 0.003 | -0.35 |

* Note: *P < 0.0125
